# A Fitness–Entropy Compensation effect set the trade-off between growth and gene expression in cell populations

**DOI:** 10.1101/2025.07.05.663304

**Authors:** Mathéo Delvenne, Vincent Vandenbroucke, Lucas Henrion, Maximilian Sehrt, Juan A. Martínez, Alizée Sloodts, Samuel Telek, Andrew Zicler, Frank Delvigne

## Abstract

We present findings on a Fitness–Entropy Compensation (FEC) mechanism which offsets the activation of gene circuits that compromise survival. It counteracts the resulting fitness reduction by increasing the diversity in gene expression among individual cells within the population. This diversity, quantified by the Shannon entropy, enables cells with lower expression levels to support the survival of the entire population. We investigated the presence of FEC in a range of synthetic and stress-related genetic circuits in continuous culture. Our results reveal that it effectively stabilizes cell populations by mitigating the detrimental trade-offs between growth and gene expression. This stabilization is due to the reduced growth rate of the induced phenotype that leads to environmental changes, decreases induction strength, and promotes escape from unfit states. These findings suggest that the FEC mechanism may be a universal strategy for stabilization in various cellular systems and set the basis for a quantitative description of the trade-off between growth and gene expression and its consequences at the population level.

## Introduction

Individual cells within a population activate specific gene circuits in response to stress conditions, often at the expense of cellular growth ^1–3^. Similarly, gene circuits employed in bioproduction can compromise cellular growth and both productivity and predictability ^4,5^. This trade-off between growth and gene expression is a crucial factor in understanding the physiological consequences of gene circuit activation, but its interpretation must be refined by accounting for the inherent variability in gene expression ^4,6–8^.

Our recent findings have highlighted the importance of fitness cost, or the growth reduction associated with the activation of a given gene circuit, in driving cell population diversification dynamics ^9^. We observed that low fitness costs result in relatively homogeneous cell population diversification profiles (**Figure 1A**), whereas high fitness costs lead to more heterogeneous diversification dynamics (**Figure 1B**) ^9^.

**Figure 1:**
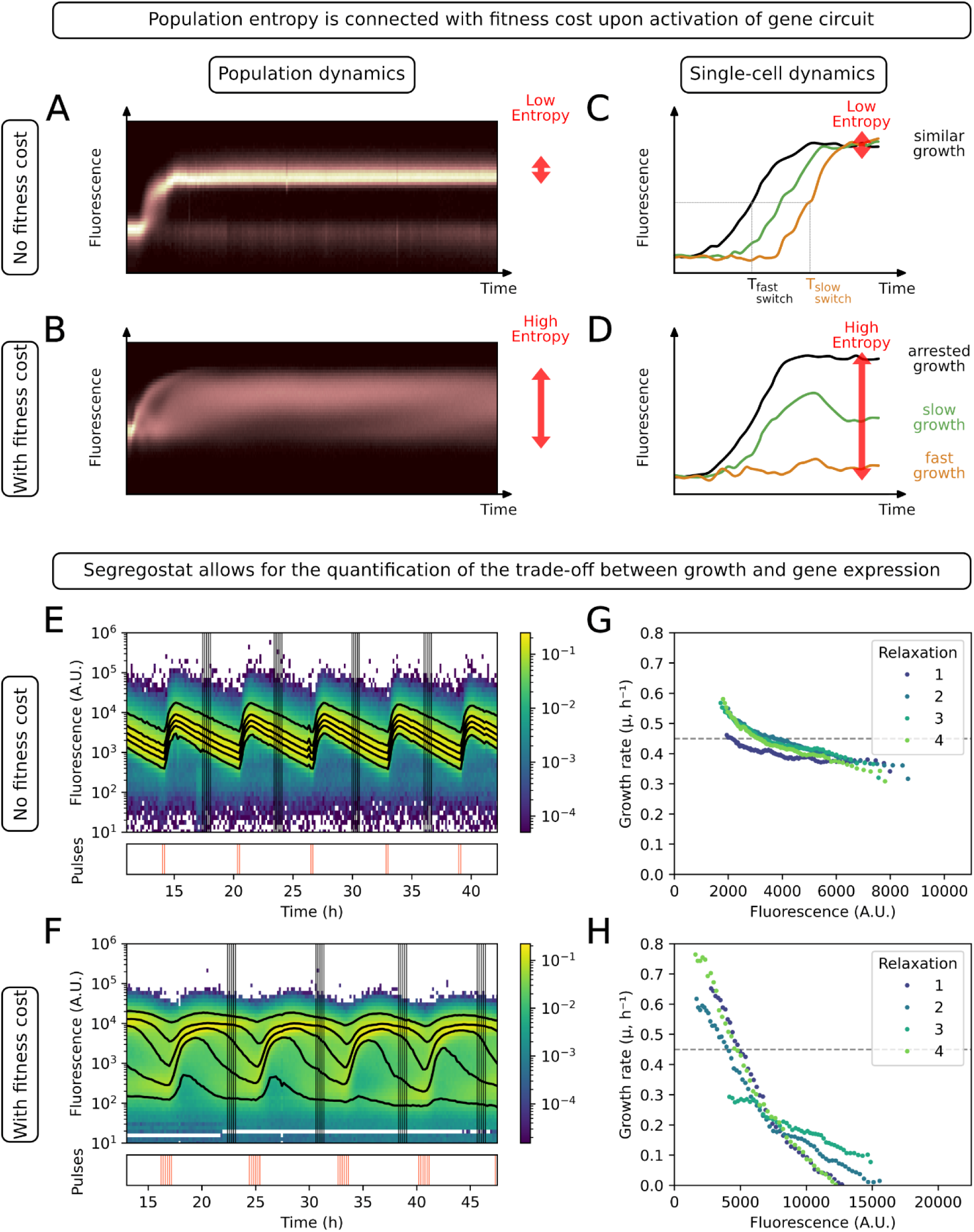
Relationship between gene expression, fitness cost, and population entropy. (A-B) Population diversification dynamics obtained based on automated flow cytometry (single-cell fluorescence distributions over time) of (A) *E. coli* P*_araB_*::GFP, without fitness cost, and (B) *S. cerevisiae* P*_glc3_*::GFP, with a fitness cost, cultivated in a glucose-limited chemostat, leading to the activation of the corresponding reporter system (adapted from Henrion *et al.* (2023)). (C-D) Schematic representation of single-cell traces for circuits with (D) or without (C) fitness cost. (E-F) Fluorescence distributions over time for (E) *E. coli* P*_araB_*::GFP and (F) *S. cerevisiae* P*_glc3_*:::GFP under Segregostat control with pulses of arabinose or glucose, respectively. Black lines represent the 5th, 25th, 50th, 75th, and 95th percentiles. Under these conditions, the fluorescence relaxation phases can be used to estimate the growth rate as a function of the fluorescence intensity. The five samples used for growth rate approximation are indicated by vertical lines. (G-H) Trade-off between gene expression (fluorescence) and growth rate for (G) *E. coli* P*_araB_*::GFP and (H) *S. cerevisiae* P*_glc3_*:::GFP based on Segregostat data, the gray dashed line indicates the dilution rate.

To quantify the degree of cell population dispersion in the activation of a given gene circuit, we utilized Shannon entropy from information theory. This metric was previously validated in various biological studies as a reliable estimator of signal transmission fidelity in gene expression ^10,11^ and it offers a robust measure of population dispersion. Notably, entropy outperforms traditional metrics such as the Fano factor and coefficient of variation, as it is mean-independent and better suited to capturing heterogeneity in multimodal distributions ^12,13^.

We hypothesized that a correlation exists between the increase in entropy observed in systems with high fitness costs and the overall fitness of the population. Previous studies have highlighted the crucial role of biological noise in enhancing the robustness of cell populations to perturbations by generating a diverse range of phenotypes adapted to various environmental conditions ^14–18^. Specifically, in the context of heterogeneous expression of burdensome genes, cells with lower expression levels may compensate for the global loss of growth at the population level. This led us to propose the concept of a Fitness–Entropy Compensation (FEC) effect, a population-level phenomenon where the loss of individual cellular fitness (e.g., due to burdensome gene expression) is offset by an increase in phenotypic diversity, measured with entropy, thereby maintaining population fitness. To investigate this effect, we first examined the T7 expression system in *Escherichia coli*, which is known for its high fitness cost. We then explored whether FEC can also occur in natural gene circuits, focusing on genes involved in the general stress response in *Saccharomyces cerevisiae* and *Escherichia coli*, as well as its potential sources.

## Results

### Segregostat reveals the trade-off between growth and gene expression

The trade-off between growth and gene expression is a fundamental aspect of cellular biology ^19,20^, yet a comprehensive framework for understanding its role at the population level remains elusive. In our previous work, we investigated this phenomenon across various cellular systems using chemostat experiments coupled with automated flow cytometry ^9^. This approach allowed us to examine the relationship between the fitness cost associated with gene circuit activation and the resulting level of heterogeneity in gene expression, quantified by entropy. Our findings revealed that non-burdensome gene circuits exhibited low population entropy (**Figure 1A**), whereas burdensome gene circuits displayed significantly higher entropy (**Figure 1B**).

This observation led us to hypothesize that growth arrest or reduction may be the driving force behind cell-to-cell differences in gene circuit activation. For non-burdensome systems, cell-to-cell variation in accumulation of fluorescent protein could be observed, likely due to natural differences in gene expression timing. However, these differences were quickly leveled out as the system reached steady state (**Figure 1C**). In contrast, burdensome gene circuits result in growth arrest and/or reduction, further amplifying the differences in fluorescent protein accumulation (**Figure 1D**). To test these hypotheses, we conducted a detailed analysis of previously acquired data from the Segregostat, a cell-machine interface that enables automated monitoring and perturbation of gene circuits based on their natural dynamics ^9,21^. This system was used to generate successive induction and relaxation phases with *E. coli* cells containing either a non-burdensome arabinose utilization reporter system or a burdensome T7 expression system (**Figures 1E & 1F**). During the relaxation phase, where fluorescence levels decrease due to protein dilution caused by cell growth and division ^22^, we investigated the relationship between fluorescence intensity (GFP signal) and growth rate (**Figures 1G & 1H, Supp. Notes 2 & 3**). In accordance with our previous fitness cost analysis ^9^, our results pointed out that the arabinose system had no significant impact on growth (**Figure 1G**), whereas the T7 system displayed a pronounced fitness cost (**Figure 1H**).

### Entropy compensates the loss of fitness upon the activation of burdensome gene circuits

The aim of this work is to determine whether entropy is linked to fitness at both the single-cell and population levels. The concept of ‘fitness’ is multifaceted and context-dependent, encompassing various levels of biological organization and timescales. In the literature, fitness can refer to individual organisms, populations, or species, and can be measured over a range of timescales, from instantaneous propensities in a defined environment to evolutionary timescales where rare events influence population dynamics ^23–27^. In this study, we consider fitness at two levels: (1) the phenotypic level, where gene induction can compromise cellular survival (e.g., growth rate in a continuous bioreactor); and (2) the population level, which comprises a diverse array of phenotypes. Furthermore, to investigate the mechanisms that enable a population to maintain its fitness over time, we consider population fitness as an instantaneous measure of the population’s adaptation to its current environment.

To analyze the potential function of entropy as a population fitness stabilizer, we performed continuous cultivation of *E. coli* with the T7 expression system on a mix of glucose and lactose. This setup yielded a complex diversification profile, characterized by bursts of diversification, where cells periodically activated the T7 system, incurred a fitness cost, and were progressively washed out (**Figures 2A & 2B**). Despite this repeated washout, the cell population exhibited a robust behavior in terms of cell density, with growth sustained by cells displaying lower induction levels (**Figure 2C**). Using automated FC data, we computed the time profiles of entropy (**Figure 2D**) and population fitness (**Figure 2E**), the latter being estimated as the mean fitness based on a trade-off curve generated from Segregostat data and fluorescence measurements (**Figure 1H, Supp. Note 3**).

**Figure 2:**
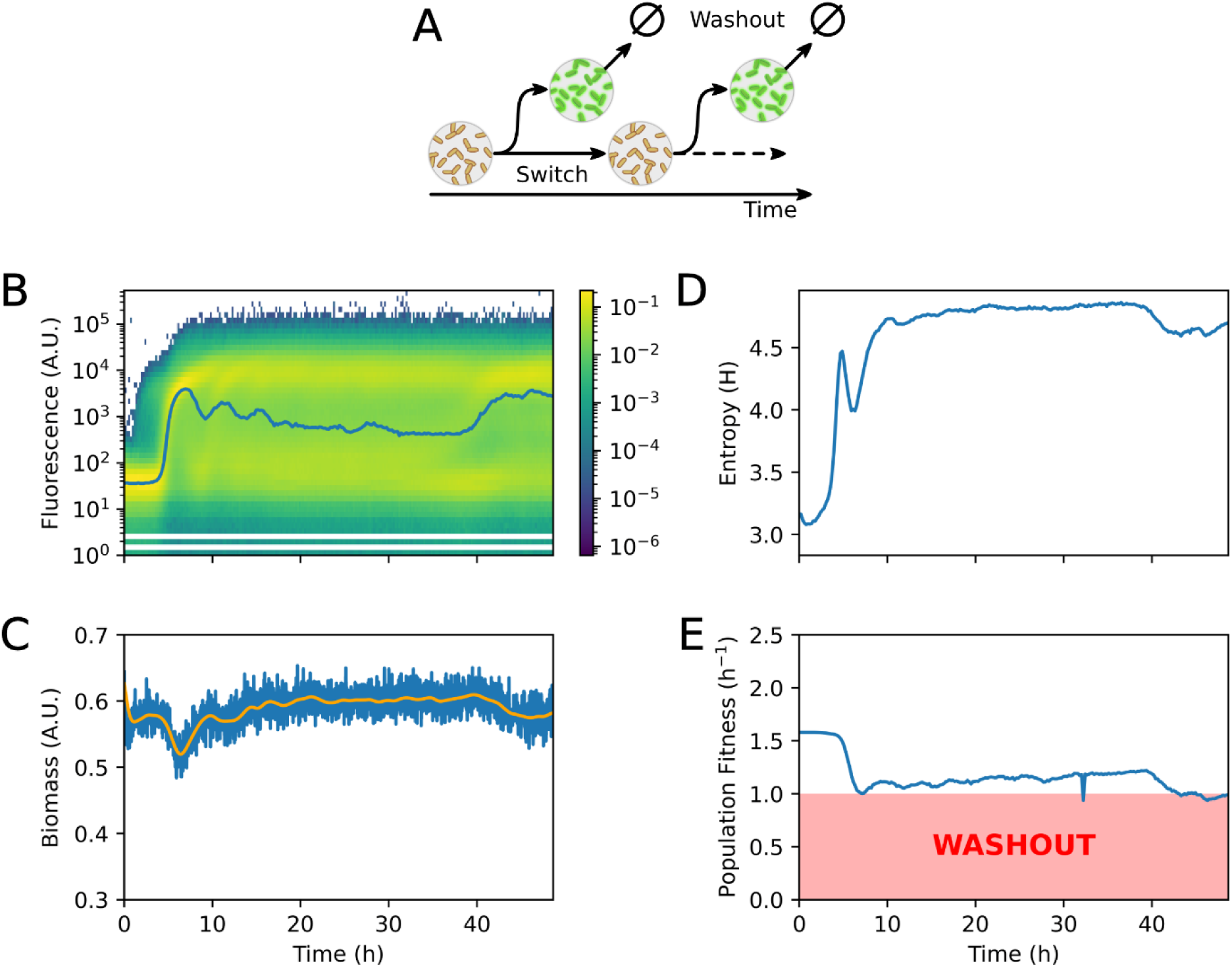
FEC for *E. coli* with a T7 expression system in continuous culture. (A) Schematic representation of the phenotypic diversification dynamics with bursts of diversification. (B) Fluorescence distributions over time determined based on automated FC (the blue line indicates the median fluorescence). (C) Biomass over time, the blue line indicates raw data and the orange line indicates smoothed data. (D) Entropy computed based on the distribution of the fluorescence over time. (E) Population fitness computed as the mean fitness based on trade-off curve generated from Segregostat data and the fluorescence from automated FC.

These data point out that the generation of entropy during the successive bursts of diversification is correlated with a global gain in fitness, keeping the global growth rate of the cell population above the dilution rate and preventing washout. Ultimately, this Fitness–Entropy Compensation (FEC) effect leads to a very stable cell density during the whole cultivation (**Figure 2C**).

A possible explanation about the origin of the FEC effect is the escape of cells from the unfit, slow-growing, states due to changes in environmental conditions (i.e., environmental escape). The bursty diversification pattern observed during chemostat cultivation suggests that the bursts of diversification may themselves trigger oscillations in the environmental conditions. Upon consumption of lactose, most of the cells are getting induced and exhibit reduced growth rate due to the fitness cost upon the activation of the T7 system, leaving out glucose in the supernatant ^28,29^. This temporary glucose increase, resulting from the feedback of cells on their environment, allows some cells to escape from the growth-limiting induction through carbon catabolite repression. Lactose and glucose concentrations are then expected to oscillate in phase opposition. This last feature will be addressed in the next section.

### FEC involves two subpopulations exhibiting opposite trade-off

In the previous section, environmental escape was advanced as a potential driver for the generation of the FEC effect. Upon induction, the T7 system exhibits a bimodal behavior with two subpopulations i.e., a slow-growing phenotype exhibiting a high induction level and a fast-growing one with a low expression level (**Figure 3A**). The co-occurrence of the two subpopulations is at the origin for the elevation of the entropy of the population, the lower subpopulations compensating for the loss of fitness of the high subpopulation. Since lactose and glucose are co-fed into the continuous cultivation device, these two subpopulations could be the result of the differential growth on a specific sugar. We validated this hypothesis by replacing glucose with xylose and observed a strong reduction in the proportion of the fast-growing, low GFP, subpopulation (**Figure 3B**). Xylose is not able to exhibit catabolite repression and is co-consumed with lactose ^30^, explaining the lack of environmental escape in this case. We then compared the measured biomass (**Figure 3C**) and entropies (**Figure 3D**) for the cultivations carried out on a mix of lactose and glucose and on a mix of lactose and xylose. When glucose is used, the entropy of the population increases during the first stage of the culture and then evolves to prevent the loss of fitness (**Figure 2C-E**, **Figure 3C & 3D**). By contrast, this is not observed when glucose is replaced by xylose. In that case, the entropy cannot be maintained, and the population collapses due to a lack of fitness (**Figure 3C & 3D**).

**Figure 3:**
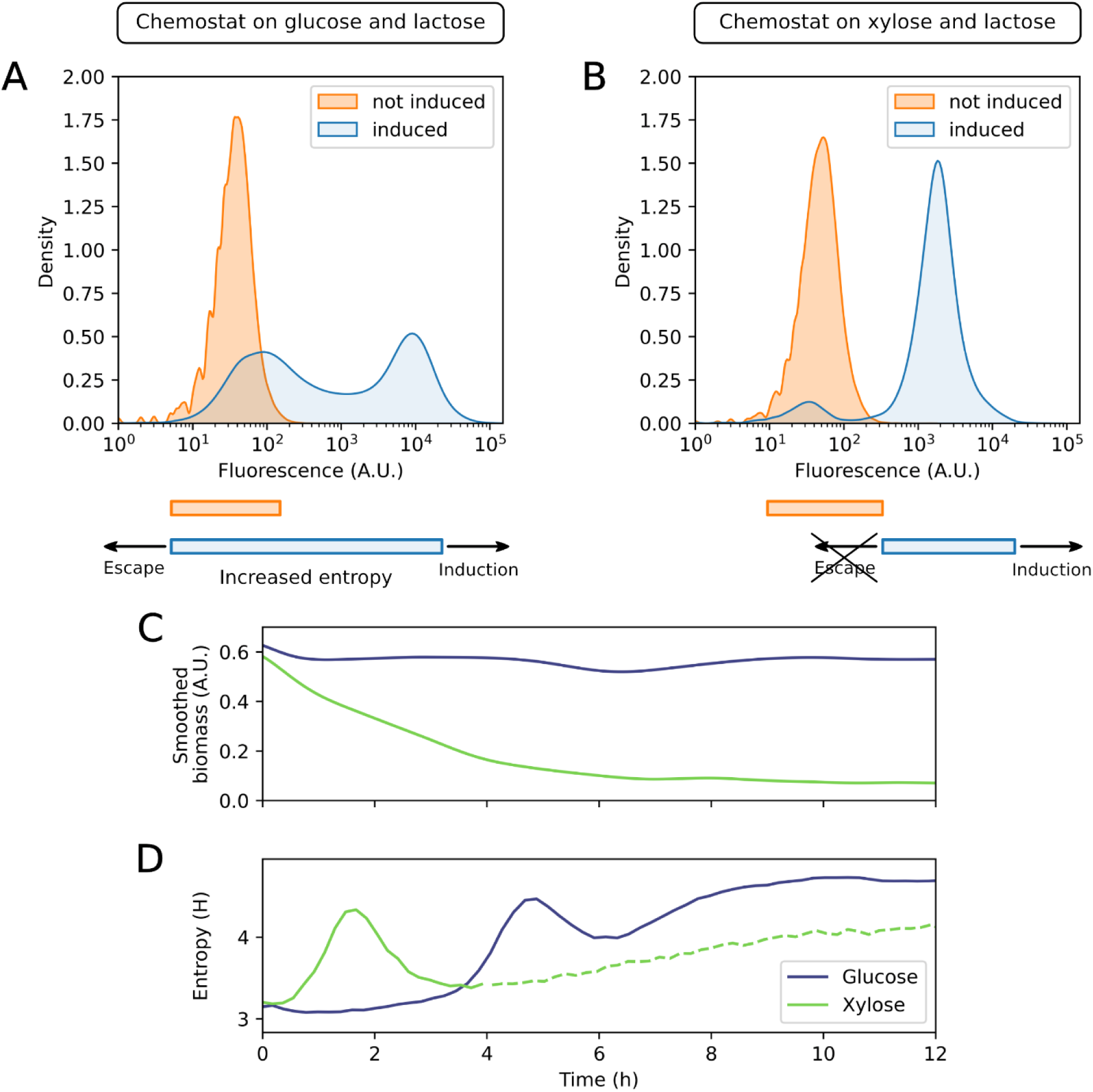
Impact of environmental escape on the heterogeneity/entropy and fitness of the population. (A-B) Fluorescence distributions of cells cultivated in chemostat on (A) glucose and lactose, and (B) xylose and lactose. Orange: before induction, blue: after induction (20 h for glucose+lactose, and 2.5 h, i.e., just before the start of washout as measured by event numbers in flow cytometry, for xylose+lactose). The schematic below the fluorescence plots illustrates the presence (A) or absence (B) of environmental escape in these conditions. (C) Smoothed biomass over time. (D) Entropy computed based on the distribution of the fluorescence over time. The dashed line indicates an increase in entropy due to a higher proportion of noise resulting from the decrease in biomass, as fewer and fewer cells are measured. In this case, no subpopulation emerges to support growth. In (C) and (D) the purple lines refer to the glucose+lactose condition, while the green lines refer to the xylose+lactose condition.

### Lack of environmental escape prevents FEC and leads to population collapse

To further explore the role of environmental escape in triggering the FEC effect, we engineered *E. coli* strains where induction is decoupled from growth, thereby isolating the impact of phenotypic switching from environmental escape. To eliminate any potential environmental escape during our experiments, we employed IPTG as an inducer. As IPTG is not metabolized by the cells, its concentration remains constant, effectively precluding any cell escape due to changes in IPTG levels.

Our negative control was a genetic toggle switch without fitness cost (**Supp. Note 4**). Upon addition of IPTG, the toggle switch is locked to a state preventing GFP production. As expected, automated FC analyses revealed that all cells exhibited a low and homogeneous GFP profile with a very low entropy and with no change in fitness at the population level.

To generate a fitness cost, the same toggle switch was used to control the *serA* gene in a serine auxotrophic *E. coli* mutant (**Figure 4A**). By placing the *serA* gene on the GFP side of the toggle switch, we created a system where IPTG induction represses GFP expression and thereby prevents serine synthesis. This resulted in a significant reduction in cellular growth rate (from 0.5 h^-1^ to 0.3 h^-1^ in microplate, **Supp. Note 4**), as the cells were unable to synthesize the essential amino acid serine. We subsequently analyzed the population dynamics of this *E. coli* strain using automated FC (**Figure 4**) and observed no FEC effect, as well as the collapse of the population upon induction of the toggle switch.

**Figure 4:**
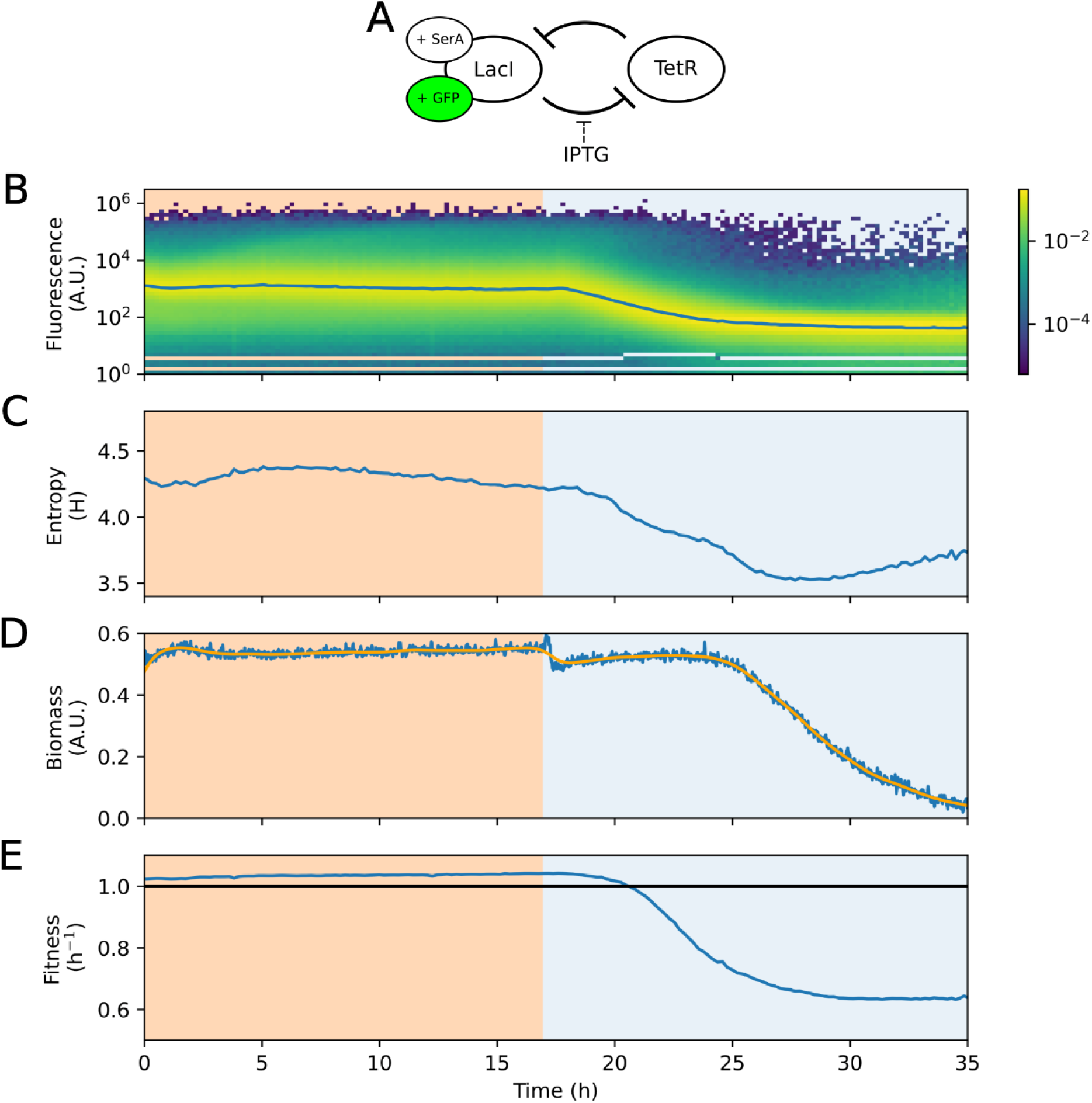
Continuous cultivation of a toggle switch controlling serine auxotrophy exhibits no environmental escape and no FEC. Upon addition of IPTG in the feed (lighter, blue background), the production of serine was inhibited. (A) Schematic representation of the toggle switch, (B) Fluorescence distributions over time as determined based on automated FC (the blue line indicates the median fluorescence). (C) Shannon entropy computed from the fluorescence distribution over time. (D) Biomass over time, the blue line indicates raw data and the orange line indicates smoothed data. (E) Population fitness computed as the mean fitness based on the fluorescence from automated FC and the independent measurement of the growth rate for each state of the switch.

In summary, our findings from two distinct gene circuits underscore the crucial role of environmental feedback in enabling cells to escape from growth-limited phenotypes through FEC.

### Segregostat and time-lapse microscopy analyses indicate that FEC is unlikely for the general stress response in *E. coli* but is expected for *S. cerevisiae*

In the previous sections, we demonstrated the occurrence of a FEC effect in synthetic gene circuits and wondered if natural gene circuits exhibited similar compensation effect. We then investigated natural stress responses in *E. coli and S. cerevisiae*. For *E. coli*, we focused on the *rpoS* regulon, which is responsible for activating various stress response mechanisms ^1,31–33^. We conducted a comprehensive search for suitable transcriptional reporters to detect the activation of the general stress response and ultimately selected the P*_ydcS_*::*GFP* system for its dynamic range and sensitivity (**Supp. Note 5**). The strong correlation between *ydcS* and *rpoS* expression confirmed its relevance as a reporter of the general stress response (**Supp. Figure S20**). For *S. cerevisiae*, we employed a previously developed P*_glc3_*::*GFP* transcriptional reporter ^9,34,35^. The *glc3* gene is associated with the general stress response in yeast and has been reported to be involved in bet-hedging ^36–38^.

To visualize the trade-off between growth and gene expression of these systems, we monitored them by time-lapse microscopy in a Microfluidic Single Cell Cultivation (MSCC) device. This system enables the cultivation of microcolonies as cell monolayers under precisely controlled environmental conditions to study cellular responses ^6,39,40^. The activation of the *ydcS* gene in *E. coli* system did not induce growth arrest in cells exhibiting high expression levels (**Figures 5A & 5B**). Instead, we observed higher global gene expression in low-nutrient conditions, accompanied by continuous growth of microcolonies. These observations were confirmed by the results of the Segregostat experiment indicating a negligible trade-off for the general stress response in *E. coli* (**Figures 5C & 5D, Supp. Note 3**). Given the significant impact of the stringent response on growth ^41^, one might expect a substantial fitness cost and corresponding increase in heterogeneity. However, *E. coli* exhibits mutual regulation between growth rate and stress response, allowing it to dynamically adjust its physiology ^1,31–33,42,43^. The general stress response is regulated by the RpoS-containing RNAP holoenzyme (EσS). The concentration of RpoS and its competition with other sigma factors for the RNA polymerase core enzyme result from complex regulation at multiple levels^2,31^. While EσS represses processes associated with rapid cell proliferation, it does not completely inhibit growth. Instead, it facilitates the expression of stress-adaptive genes that support survival and limited metabolic activity. Notably, in certain contexts, RpoS is required for maintaining slow growth by sustaining the expression of central metabolism genes ^31,44^. We carried out similar analyses on the P*_glc3_*::*GFP* system in yeast at high and low nutrient conditions ^9,35^. In both cases, cells expressing the *glc3* reporter exhibited growth arrest, which ultimately halted the expansion of the entire microcolony in the case with low glucose concentrations (**Figures 5C & 5D**). This drastic growth arrest was confirmed based on Segregostat experiments (**Figure 5G**), with a trade-off curve pointing out a substantial reduction in growth rate upon full activation of the reporter system (**Figure 5H, Supp. Note 3**). This aligns with previous reports showing that, unlike *E. coli*’s, *S. cerevisiae*’s investment in stress resistance appears incompatible with proliferation, switching between growing and non-growing states ^3,36,45^.

**Figure 5:**
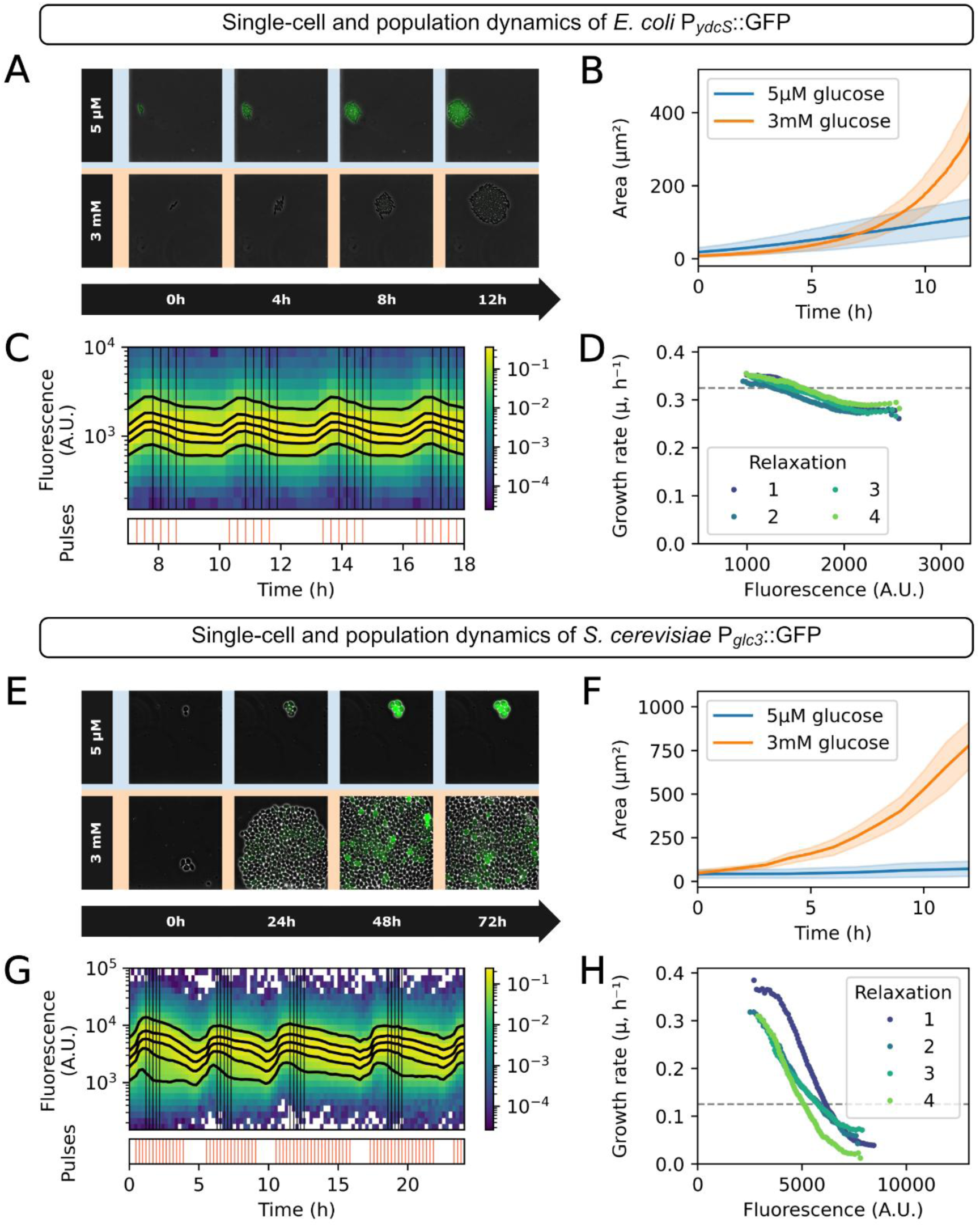
Single cell and population dynamics in MSCC and in Segregostat, for *E. coli* P*_ydcS_*::GFP and *S. cerevisiae* P*_glc3_*::GFP. (A & E) Time-lapse fluorescence microscopy images of colonies grown in mineral medium with 5 µM (top, blue) or 3 mM (bottom, orange) glucose concentration. (B & F) Colony growth in microfluidic chambers, quantified as the total cell area over time (for (B) *E.* coli, n=3 for both 5 µM and 3 mM glucose; for (F) *S. cerevisiae*, n=4 for 5 µM glucose and n=3 for 3 mM glucose). (C & G) Fluorescence distributions over time under Segregostat control with pulses of glucose. Black lines represent the 5th, 25th, 50th, 75th, and 95th percentiles. The five samples used for growth rate approximation are indicated by vertical black lines. (D & H) Trade-off between gene expression (fluorescence) and growth rate based on Segregostat data, the gray dashed line indicates the dilution rate.

The lack of a clear trade-off upon activation of the general stress response in *E. coli* suggests that the population is homogeneous with individual cells growing at a similar rate during the relaxation phase. It implies that no FEC effect is anticipated for this system, in contrast to the general stress response in yeast, where a significant trade-off was observed. This prediction will be further explored in the next section, based on experiments carried out in continuous cultures.

### FEC is also involved in natural stress response systems exhibiting a marked trade-off between gene expression and growth

To investigate the potential occurrence of FEC in natural gene circuits involved in the general stress response, we cultured our reporter strains in chemostats and monitored them using automated flow cytometry. Cultivations were carried out at a high dilution rate (non-stressful control condition) and a low dilution rate (nutrient-limited, stress-inducing condition) for comparison.

For the *E. coli* P*_ydcS_*::*GFP* reporter, switching the chemostat from a high (D = 0.6 h^-1^) to a low dilution rate (D = 0.1 h^-1^) led to a strong and homogeneous activation of the reporter system (**Figures 6A & 6E**). The entire population transitioned from the non-stressed to the stressed phenotype, with the fluorescence distribution shifting uniformly from low to high GFP levels. The two corresponding distributions show minimal overlap, indicating that *E. coli* precisely modulates the expression of stress-related genes in proportion to stress intensity, functioning like a rheostat. ^46^. Furthermore, entropy did not increase upon switching to the stressed state; it may even be slightly reduced (**Figures 6B & 6E**). Notably, fitness remained unaffected (**Figure 6D**). Since the general stress response in *E. coli* does not substantially impact growth under these conditions (even if it is reduced, the growth rate remains above the dilution rate), the FEC mechanism does not appear to be involved in this response.

**Figure 6:**
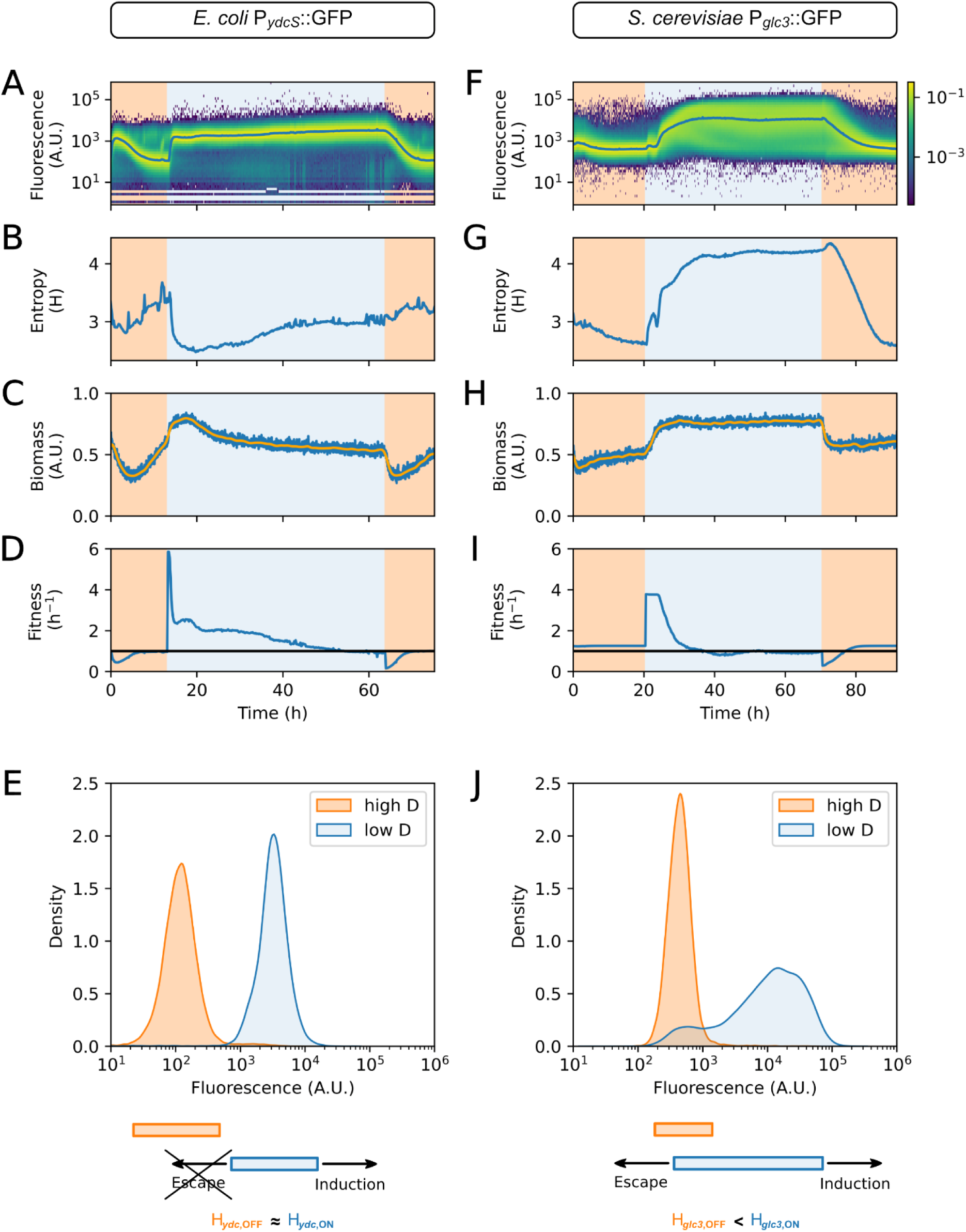
Single cell and biomass analysis of (A-E) *E. coli* P*_ydcS_*::GFP and (F-J) *S. cerevisiae* P*_glc3_*::GFP cultivated in continuous. cultures carried out at high (darker, orange background, D = 0.6 h^-1^ for *E. coli* and D = 0.3 h^-1^ for *S. cerevisiae*) and low (lighter, blue background, D = 0.1 h^-1^ for both strains) dilution rates. (A & F) Fluorescence distributions over time determined based on automated FC (the blue line indicates the median fluorescence). (B & G) Shannon entropy computed from the corresponding fluorescence distribution over time. (C & H) Biomass over time (the blue line indicates raw data and the orange line indicates smoothed data). (D & I) Population fitness computed as the mean fitness based on trade-off curve generated from Segregostat data and the fluorescence from FC. (E & J) Fluorescence distributions of samples representative of cells cultivated in chemostat at high (t = 75 h for *E. coli* and t = 90 h for *S. cerevisiae*) and low (t = 63 h for *E. coli* and t = 70 h for *S. cerevisiae*) dilution rates. The schematic below the fluorescence plots illustrates the absence (E) or presence (J) of environmental escape.

For the *glc3* system in yeast, time-lapse microscopy and Segregostat experiments revealed that strong activation can reduce growth below the dilution rate, leading to growth arrest when fully activated (**Figures 5E-H**). Accordingly, we might expect population washout upon switching to the low dilution rate (D = 0.1 h^-1^), but this was not the case. In stark contrast to *E. coli*, *S. cerevisiae* displayed a stress response consistent with the “noisy switch model”, characterized by overlapping gene expression distributions across varying stress intensities ^46^ (**Figures 6F & 6J**). At low dilution rates, the population split into two distinct subpopulations: one continued to elevate its stress response, while the other reverted toward pre-stress expression levels. This resulted in a bimodal fluorescence distribution and a significant increase in entropy (**Figure 6G**). Notably, population fitness did not decrease (**Figure 6I**), and biomass concentration remained stable (**Figure 6H**). The observed increase in entropy may therefore reflect a functional role at the population level, enabling partial escape from the slow-or non-growing stressed state and thereby preventing washout in continuous cultivation. This is consistent with the hypothesis that increased entropy compensates for the growth reduction associated with the activation of the stress reporter.

These observations confirm the fact that considerable reduction in growth rate upon activation of the target gene circuit is a crucial component of the FEC mechanism. The second condition previously reported for the generation of the FEC effect observed for the *glc3* system in yeast is the occurrence of an environmental escape mechanism. A possible explanation for the partial relaxation of cells from the stressed to the non-stressed state in low-dilution rate chemostats is that stressed cells exhibit reduced metabolic activity and tend to shift toward a more quiescent state, thereby decreasing substrate consumption ^3,36,45^. Fully stressed cells may even cease growth and nutrient uptake entirely. This reduced consumption enables cells with lower stress-related gene expression to benefit from a less depleted environment, allowing them to escape the stressed state via an environmental feedback mechanism.

Moreover, when cultivated in the MSCC device under low glucose concentrations, strongly activated cells remained locked in the stresses state, with no observable decrease in fluorescence (**Figures 5E & 5F, Supp. Figure S23**). In microfluidics, cell physiology is decoupled from environmental conditions, thereby eliminating environmental feedback. Residual glucose concentrations ([Glu_res]) were measured in continuous cultures across various dilution rates and compared to those in the MSCC device (**Supp. Note 7**). At low dilution rates (D < 0.3 h⁻¹), [Glu_res] remained very low around 0.1 mM — the threshold concentration at which the entire population exhibited growth arrest in the MSCC device (**Supp. Figures S23B**). This supports the hypothesis that transient environmental conditions enable cells to escape the stressed state.

## Discussion

Gene expression noise and phenotypic switching enables microbial populations to adapt and thrive in dynamic environments ^14–16,47–49^. To date, most studies have treated phenotypic diversification both as a consequence of intrinsic gene expression noise and as an adaptive strategy for coping with fluctuating environments. However, with a few notable exceptions ^16,50,51^, the impact of phenotypic changes on population composition has received limited attention. This knowledge gap is critical, as fitness variation among phenotypes inevitably reshapes population structure through intra-population competition. Our study seeks to address it by laying the groundwork for a robust experimental framework to better understand these dynamics.

A well-documented functional consequence of noise in gene expression is bet hedging ^14–16,49^, a strategy microbial populations use to enhance survival in unpredictable environments. This mechanism leverages noise to generate a small subset of phenotypes that, while suboptimal under current conditions, may prove advantageous upon environmental shifts. These alternative phenotypes typically represent a minor fraction of the population, as they are usually outcompeted by the dominant, better-adapted phenotype ^16^. But what happens when the dominant phenotype becomes maladapted to its environment? In such cases, the population would be expected to die out. However, several studies revealed a surprising resilience, with populations surviving conditions that would normally be lethal. Besides this work, a prominent example is bacterial persistence, were a resistant phenotype temporarily outcompetes sensitive cells even without mutations ^52^. This unexpected survival can be explained by a Fitness– Entropy Compensation (FEC) mechanism. According to this mechanism, when the dominant phenotype fails to persist, a minority of cells that stochastically escape this state — via variability in gene expression or repression — can proliferate and repopulate the culture. This competitive dynamic is essential, as it prevents any single phenotype from dominating the population. Instead, it promotes an equilibrium among phenotypes, resulting in increased heterogeneity (**Figure 6**). This heterogeneity is quantitatively captured by an increase in population entropy (as shown mathematically in **Supp. Note 1**).

The emergence of alternative phenotypes is facilitated by an environmental escape mechanism. Under our chemostat conditions, both *S. cerevisiae* under glucose starvation and *E. coli* BL21 grown on a mixture of glucose and lactose adopt phenotypes that are maladapted to survival. In *S. cerevisiae*, glucose limitation induces a non-growing, stress-related phenotype, while in *E. coli* BL21, lactose consumption triggers the activation of the burdensome T7 gene circuit, significantly reducing growth rate. In both cases, the decline of the induced, unfit subpopulation leads to a rise in environmental glucose levels. This, in turn, inhibits further induction of the maladapted phenotype — by relieving stress in yeast and via catabolite repression in BL21. As a result, the microbial population naturally adjusts its environment to a glucose concentration that permits sufficient gene expression noise for FEC to occur. This feedback mechanism is likely to arise in any context where the decline of the microbial population reduces induction strength.

The analysis of the FEC mechanism was enabled by quantifying the trade-off between growth rate and gene expression. In some systems, such as the growth-limiting toggle switch, it is possible to isolate and measure two clearly distinct phenotypes. However, biological systems are rarely so clear-cut. Microfluidics is the alternative option to map fitness to phenotype, but while it provides valuable temporal resolution, it is labor-intensive and does not accurately replicate the growth conditions of our systems of interest. By contrast, our approach using automated FC offers a direct and efficient means to assess growth rates within bioreactor environments. We propose that this technique represents a critical step towards a more robust understanding of bacterial population dynamics. Notably, we were only able to quantify a single trait at once, and an important question we did not address is whether additional heterogeneity, such as variations in transporter expression or metabolic regulation, could influence which cells escape induction. This could be investigated through the use of multiple reporters ^53,54^ or via omics approaches on subpopulation or even at the single cell level ^55^.

Overall, our findings seem to have far-reaching implications across biological systems. We measured FEC in chemostats, but the phenomenon is applicable to microbial population dynamics anywhere a fitness can be described, whether a bioprocess, an antibiotics resistance study, or a biofilm layer ^52,56^. In the laboratory, where FEC is relevant, applying results from one setting to another demands careful consideration because fitness and environmental escape are context dependent. When studying diversity itself, FEC alters how much gene expression noise is required for a species to effectively hedge its bets against a fluctuating environment and could affect our understanding of both the evolutionary path of microbes and the expression of recombinant protein genes in industrial settings.

## Material and Methods

### Code and data availability

All relevant code is available at https://gitlab.uliege.be/mipi/published-software/2024-fitnessentropycompensation. All supplementary data are available at doi.org/10.5281/zenodo.14273182, besides the raw time-lapse microscopy files that can be made available on request due to their large size. *<u>and plasmids</u>*

A strain of *E. coli* W3110 with a plasmid from the Zaslaver collection (*P_araB_*::GFPmut2 with kanamycin resistance ^57^) was used as an example of arabinose utilisation system with low fitness cost. The data acquired with this strain were already published ^9^.

The *E. coli* BL21(DE3) strain, one of the most important industrial production backbone, carrying the pET28:GFP plasmid (<u>Addgene #60733</u>)^58^ was used to investigate the burdensome T7 expression system. Part of the data acquired with this strain were already published ^9^.

For the toggle switch study, the control strain without growth change is *E. coli* MG1655 Δ*lacAYZ* and Δ*lacI* with the toggle-switch plasmid pECJ3 (<u>Addgene #75465</u>)^59^. To obtain the strain with auxotrophy to the plasmid, the Δ*serA* modification was added to the genome and the *serA* gene was added to pECJ3 between the promoter for GFP and the GFP (P_LtetO-1_::*serA*::GFP) with the same RBS as GFP, so that the latter is a direct reporter for serA content. The genomic modifications were performed using the CRISPR-cas9 enhanced lambda red phage mediated homologous recombination method described by Jiang et al. (2015). *lacAYZ* and *lacI*, being contiguous, were removed in one step and *lacI* was used as target, which also eliminates any hypothetical copy of *lacI* in the episome. The primers used are available in **Supp. Table 2**. The modified plasmid was produced through PCR and NEBuilder^®^ HiFi DNA Assembly; the plasmid map is available in the supplementary data. Each phenotype of the strain with auxotrophy to its plasmid was measured in Biolector, a microfermentation system enabling online monitoring of biomass and fluorescence in microtiter plates. It showed that the growing side has a growth rate >0.5 h^-1^, while the non-growing side has a growth rate between 0.25 and 0.3 h^-1^, slow enough to be washed out of the bioreactor in continuous culture, but not zero, likely due to leaky repression of serine (**Supp. Note 4**).

For the study of the general stress response of *Escherichia coli* in chemostat, we utilized the *E. coli* MG1655 P*_ydcS_*::GFPmut2 strain from the Zaslaver collection ^57^. This strain carries a plasmid containing a transcriptional reporter for P*_ydcS_* and confers kanamycin resistance. The same plasmid was introduced into the *E. coli* MG1655 *P_rpoS_*::CyOFP1 strain to generate the double-reporter strain used in microfluidics. *E. coli* MG1655 *P_rpoS_*::CyOFP1 was constructed using the same CRISP-cas9 system as above. The CyOFP1 gene was inserted downstream of the rpoS gene with a synthetic ribosomal binding site (RBS) 5’-AAAGAGGAGAAA-3’ ^54,61^. The cassette containing the coding sequence (CDS) of the CyOFP1 protein has been obtained by pcr overlap of the CyOFP1 sequence (obtained from the pNCS-CyOFP plasmid (<u>Addgene #74278</u>) and two homologous regions of 400bp around the site of insertion. Tailed primers have been used to include the synthetic RBS in the cassette. The primers used are described in **Supp Table 1.**

For the study of the general stress response of *Saccharomyces cerevisiae* in chemostat and in microfluidics, we utilized S. cerevisiae CEN.PK 113-7D strain with the chromosomal integration of a reporter cassette *P_glc3_*::eGFP ^34,62^.

### Growth media

Unless otherwise specified (e.g., regarding the carbon source and its concentration), the following media were used.

*E. coli* strains have been cultivated in a mineral medium containing (in g/l), K_2_HPO_4_: 14.6, NaH_2_PO_4_ ⋅ 2H_2_O: 3.6, Na_2_SO_4_: 2, (NH_4_)_2_SO_4_: 2.47, NH_4_Cl: 0.5, (NH_4_)_2_−H−citrate: 1, glucose: 5, thiamine: 0.01. The medium is supplemented with a trace element solution totaling 11 ml/l assembled from the following solutions (in g/l), 3/11 of FeCl_3_ ⋅ 6H_2_O: 16.7, 3/11 of EDTA: 20.1, 2/11 of MgSO_4_: 120, and 3/11 of a metallic trace element solution. The metallic trace element solution contains (in g/L): CaCl_2_ ⋅ 2H_2_O: 0.74, ZnSO_4_ ⋅ 7H_2_O: 0.18, MnSO_4_ ⋅ H_2_O: 0.1, CuSO_4_ ⋅ 5H_2_O: 0.1, CoSO_4_ ⋅ 7H_2_O: 0.21. For plasmid maintenance, it has been supplemented with kanamycin (50 mg/l). The trace element solution, the thiamine and the antibiotic were filter-sterilized (0.22 µm) before supplementation of the other component that were heat-sterilized (at 120°C).

*S. cerevisiae* has been cultivated in a mineral medium based on the recipe of Verduyn et al., (1992). It contains, per liter, (NH_4_)_2_SO_4_: 5 g, KH_2_PO_4_: 3 g, MgSO_4_ ⋅ 7H_2_O: 0.5 g, EDTA: 95.55 mg, ZnSO_4_ · 7H_2_O: 22.5 mg, MnCl_2_ · 4H_2_O: 5 mg, CoCl_2_ · 6H_2_O: 1.5 mg, CuSO_4_ · 5H_2_O: 1.5 mg, Na_2_MoO_4_ · 2H_2_O: 2 mg, CaCl_2_ · 2H_2_O: 22.5 mg, FeSO_4_ · 7H_2_O: 15 mg, H_3_BO_3_: 5 mg, KI: 0.5 mg and glucose: 7.5 g. After heat sterilization (at 120°C) of those components, the filter-sterilized (0.22 µm) vitamins were added. The final medium contains per liter, D-biotin: 0.05 mg, calcium pantothenate: 1 mg, nicotinic acid: 1 mg, myo-inositol: 25 mg, thiamine HCl: 1 mg, pyridoxine HCl: 1 mg and para-aminobenzoic acid: 0.2 mg.

### Continuous cultivations

Continuous cultivations were performed in either Biostat B-Twin (Sartorius, Göttingen, Allemagne) or in Bionet F1 bioreactor (Bionet, Murcia, Spain), with a working volume of 1 l, at an agitation speed of 1000 rpm, with an aeration of 1 VVM. The temperature control was set either at 37 °C (*E. coli*) or 30 °C (*S. cerevisiae*) while the pH control was set either at 7 (*E. coli*) or 5 (*S. cerevisiae*). Bioreactor cultivations started with a batch phase lasting until full consumption of the carbon source (marked by an increase in oxygen concentrations) or after ∼15 h for yeast. During the chemostat experiment, glucose was fed alone or with lactose for experiment with *E. coli* BL21 strain. During Segregostat experiments, glucose was the sole carbon source in the feed. The feedback control loop of the Segregostat (implemented via a custom MATLAB script exploiting flow cytometry data, the acquisition of which is detailed below) triggered a pump to deliver pulses of the activator. For inducible systems, the actuator was the corresponding inducer (i.e., arabinose for *E. coli* P*_araB_*::GFP and lactose for *E. coli* BL21). For systems with stress response reporters, glucose was pulsed instead. The control was based on a threshold of fluorescence and the pulse was activated when more than 50% of the population was below this threshold. The specific operating conditions of the continuous cultures performed in lab-scale bioreactors are described below and summarized in **Supp. Table 3**.

*E. coli* P*_araB_*::GFP and *E. coli* BL21 were precultured in 1 L baffled flask (with 100 ml of medium) overnight at 37 °C at a shaking speed of 150 rpm. They were used to inoculate the reactor with an initial OD_600_ of ∼0.5. Continuous cultures were performed at a dilution rate of 0.45 h^-1^.

The toggle-switch strains were grown only in the same defined media as described above as a change in medium conditions would induce an unpredictable lag phase for the Δ*serA* strain. Glycerol stocks were prepared from a culture in shake flask with 10 g/L glucose medium and an OD above 5 (which is the maximal cell density reached with 5 g/L glucose). The pre-cultures (40 ml of medium in 500 ml flasks) were then inoculated directly from glycerol stock solutions and incubated for 18 to 24 h at 37° at a shaking speed of 150 rpm. The continuous cultures were performed with a dilution rate of 0.5 h^-1^. For both toggle-switch strains, the continuous culture first ran for 12 h without inducer. The feed was then replaced by a feed supplemented with 0.1 mM IPTG to induce the culture. For the strains with the unmodified toggle-switch, the continuous culture was run for 32 h with IPTG then stopped. For the toggle with growth control, the IPTG feed was kept for approx. 20 h.

Strains with stress-related genes reporters were pre-cultivated in two steps to avoid the induction of the reporter. Both steps were performed in 500 ml or 1 L flasks (with respectively 50 or 100 ml of the corresponding medium), either at 37 °C (*E. coli*) or 30 °C (*S. cerevisiae*) at a shaking speed of 150 rpm. The first preculture was inoculated either from glycerol stock or with one single colony from a LB (supplemented with 50 mg/L kanamycin, for *E. coli*) or a YPD (*S. cerevisiae*) agar plate, and was incubated overnight. The second preculture was inoculated with cell suspension from the first preculture, to get an initial OD_600_ ≈ 0.5, and was incubated until it reached enough biomass to inoculate the bioreactor (∼3 h for *E. coli*, ∼6 h for *S. cerevisiae*). The batch phase was initiated with an OD_600_ ≈ 0.1 (or 0.5 for experiments already published in previous paper ^9^). The chemostat conditions alternated between high (D = 0.6 h^-1^ for *E. coli* or D = 0.3 h^-1^ for *S. cerevisiae*) and low (D = 0.1 h^-1^) dilution rates, with each condition maintained for a duration of at least 5 residence times. The Segregostat experiment of E. coli was performed at an intermediary dilution rate (D = 0.3 h^-1^). For Segregostat experiments with yeast, a low dilution rate (D = 0.1 h^-1^) was applied and, the glucose concentration of the feed was 5 g/L.

Dissolved oxygen and pH were monitored in all experiments. Excepted for experiment previously published ^9^, biomass was monitored with Hamilton Dencytee RS485 probes. All cultures were also monitored at the single-cell level using automated flow-cytometry. The automated flow-cytometry system is the combination of an in-house sampling device (part of the Segregostat ^9,21^) and a benchtop flow cytometer (BD Accuri C6 or BD Accuri C6+). Every ∼12 min, it takes a sample from the bioreactor and dilutes it with PBS before analysis. For data acquired with the BD Accuri C6 flow cytometer, the fluidics flow rate was set to custom (24 µL/min, 8 µm core) and a threshold was set either to 20 000 (*E. coli*) or 80 000 (*S. cerevisiae*) units of FSC-H. For the data acquired with the BD Accuri C6+ flow cytometer, the fluidics flow rate was set to medium (35 µL/min, 16 µm core) and a threshold was set either to 15 000 (*E. coli*) or 80 000 (*S. cerevisiae*) units of FSC-H. Under these conditions, about 40 000 cells were analyzed per sample. The FL1-A channel (exc. 488 nm, em. 533/30 nm or em. 510/10 nm for the toggle-switch) was used for GFP quantification.

### Continuous cultivations: data treatment

To allow comparison of replicates acquired with different flow cytometers, we applied transformations on the data acquired with the BD Accuri C6 (**Supp. Note 2**). Flow cytometry data were cleaned by removing zeros and doublets. To compute the entropy, 50 fixed bins in log scale between 1 and 10⁶ a.u. were used. Biomass data was smoothed using a Butterworth lowpass filter with cutoff frequency 0.4 h^-1^.

The population fitness (or expected fitness) was computed based on flow-cytometry data as the mean growth rate of single cells in the population divided by the dilution rate. The single cell growth rate was inferred using functions linking fluorescence and growth rate. For most of our systems, this function was determined by exploiting the fluorescence relaxation phase in Segregostat (**Supp. Note 3**). Regarding *E. coli* P*_ydcS_*::GFP, no trade-off was observed between general stress expression and growth rate, making this computation less relevant. However, given the homogeneity of the population, an approximation of the function linking single-cell fluorescence and growth rate was obtained using chemostat data acquired at various dilution rates (**Supp. Figure S14**). For the toggle switch system, we considered two phenotypes: GFP^+^ (able to produce serine) and GFP^-^ (without serine production) separated by a threshold of 100 fluorescence a.u.. In this case, the population fitness was calculated as a weighted average of the growth rates of the two subpopulations based on their relative abundances divided by the dilution rate:

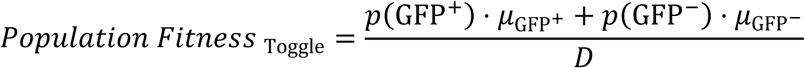

Where D is the dilution rate and p(GFP+) and p(GFP−) represent the proportions of GFP⁺ and GFP⁻ cells in the population. Their respective growth rates µ_GFP+_ = 0.531 +-0.014 (std) h^-1^ and µ_GFP-_ = 0.295 +/- 0.010 (std) h^-1^ were determined during the toggle-switch phenotypic characterization (**Supp. Note 4**).

### Microfluidic cultivations and time lapse microscopy

Cells precultures were performed in the same way as those for the lab-scale bioreactor, but in smaller flasks (100 ml with 10 ml of the corresponding medium).

The media used are the same as described above, but with modified glucose concentrations (5 µM for stressful conditions and 3 mM for non-stressful conditions).

Cells have been cultivated in microfluidic chips provided by Alexander Grünberger’s lab. They are PDMS-glass based microfluidic single-cell cultivation (MSCC) device featuring mono-layer growth (chambers sizes: 80 μm x 90 μm x ∼850 nm for *S. cerevisiae* and 80 μm x 90 μm x ∼700 nm for *E. coli*) ^40,64,65^. The experiments on *S.cerevisiae*, were already published ^9,35^. They were reprocessed to calculate colonies areas using the same method as for *E. coli* (see below). At least 3 cultivation chambers were selected manually for each glucose concentration condition. The settings of the experiments carried out with *E. coli* are slightly different. The temperature was set at 37°C. The chambers were inoculated with a few cells by flushing the device with a cell suspension (OD_600_ ≈ 0.1). Three biological replicates have been performed for each glucose concentration condition. For two of them, one chamber was analyzed while 3 chambers (technical replicate) were analyzed in the last biological replicate (for a total of 5 chambers for each glucose concentration condition).

Microscopy images of *E. coli* were acquired using a Nikon Eclipse Ti2-E inverted automated epifluorescence microscope (Nikon Eclipse Ti2-E, Nikon France, France) equipped with aDS-Qi2 camera (Nikon camera DSQi2, Nikon France, France), a 100× oil objective (CFI P-Apo DM Lambda 100× Oil (Ph3), Nikon France, France). The GFP-3035D cube (excitation filter: 472/30 nm, dichroic mirror: 495 nm, emission filter: 520/35 nm, Nikon France, Nikon) and the TxRed-4040C cube (excitation filter: 562/40 nm, dichroic mirror: 593 nm, emission filter: 624/40 nm, Nikon France, Nikon) were used to measure GFP and CyOFP1, respectively. The phase contrast images were recorded with an exposure time of 300 ms and an illuminator’s intensity of 30 %. The GFP images were recorded with an exposure time of 500 ms and an illuminator’s intensity of 2 % (SOLA SE II, Lumencor, USA). The CyOFP1 images were recorded with an exposure time of 800 ms and an illuminator’s intensity of 50 % (SOLA SE II, Lumencor, USA). Phase contrast images were acquired every 5 min while both GFP and CyOFP1 fluorescence were measured every 20 min. The optical parameters and the timelapse were managed with the NIS-Elements Imaging Software (Nikon NIS Elements AR software package, Nikon France, France).

The cell segmentation of the images of *S. cerevisiae* was performed using the Matlab algorithm of Wood and Doncic ^66^. Microscopy images of *E. coli* were analyzed as follows. First, the images were exported in ‘.tiff’ files using the NIS-Elements Imaging Software (Nikon NIS Elements AR software package, Nikon France, France). Then, they were preprocessed (normalization, stabilization, cropping, etc.) using a custom-made Python pipeline and Fiji/ImageJ. The cell segmentation was performed on phase contrast images using a custom-made Python code based on cellpose library (version 0.7.1) ^67,68^ with the trained model of Dimalis (https://github.com/Helena-todd/Dimalis, commits 26 Jul 2023). Masks images of segmented cells were corrected manually using Fiji/ImageJ before tracking using the Fiji/ImageJ plugin TrackMate (version 7.11.1) ^69,70^ with the “Label image detector” and the “Overlap tracker” options (precise IuO calculation, min. IuO = 0.1 and scale factor = 1.1). Singles-cells data and lineages were then exported as tables and further analyzed. For instance, different approximations of the growth rates have been performed. The one used in **Supp. Figure S21** is based on the increased area during the life of single cells (until division) considering an exponential growth. This approximation is computed based on filtered area (to decrease the noise due to segmentation artefacts between time frames).

## Acknowledgements

MD is supported by a PhD grant provided by the FRS–FNRS and the H2020 program of the European commission (Era-Net Aquatic Pollutant project ARENA). VV and LH are supported by a FRIA grant provided by the “Fonds de la Recherche Scientifique” FRS– FNRS from the Walloon region of Belgium. JAM is supported by a post-doctoral grant provided by the Service Public de Wallonie (SPW) and the H2020 program of the European commission (Era-Cobiotech project Contibio). FD received funding from a research grant provided by the Service Public de Wallonie (SPW) and the H2020 program of the European commission (Era-Cobiotech project ComRaDes). We would like to warmly thank Laurie Josselin for her valuable feedback and suggestions during the preparation of this manuscript and all the FD Team for their support, discussions and ideas.

