## Supplementary material for "A Fitness–Entropy Compensation effect set the trade-off between growth and gene expression in cell populations"

Supplementary materials for
### Fitness Entropy compensation

First version

Mathéo Delvenne[a,1], Vincent Vandenbroucke[a,1], Lucas Henrion[a], Maximilian Sehrt[a], Juan Andres Martinez[a], Alizée Sloodts[a], Samuel Telek[a], Andrew Zicler[a], and Frank Delvigne[a,2]

[1] V.V. and M.D. contributed equally to this work.

[a] Terra Research and Teaching Centre, Microbial Processes and Interactions (MiPI), Gembloux Agro-Bio Tech, University of Liège, Gembloux, Belgium

4th July 2025

**Supplementary Note 2.**
**Calibration of the automated flow cytometry** **11**

**Supplementary Note 3.**
**Determination of the trade-off curves based on Segregostat experiments** **13**

**Supplementary Note 6.**
**Time-lapse microscopy in a Microfluidic Single Cell Cultivation (MSCC) device** **29**

**Supplementary Note 7.**
**Cultivations of *S. cerevisiae* $P_{glc3}$::GFP at various dilution rates** **31**

**8.**
**Tables** **32**

8.1. Strains and plasmids . . . . . . . . . . 32
8.2. Lab-scale bioreactors cultivations . . . . . . . . . . 35

**Supplementary References** **36**

### Supplementary Note 1. A theoretical analysis of the Fitness-Entropy compensation mechanism

In this note, we first introduce the concept of Shannon entropy and its application to biological measurements (this information is the foundation of information theory and can be found, for most of it, for example, in [1]). Then, starting in section 1.1., we go into the details of the measurement and interpretation of entropy in the context of biological measurements, why entropy should increase mechanically as a result of the competition in simplified and idealised cases (section 1.3.), and finish with a more detailed reasoning that can be applied to realistic cases such as the ones we measured (section 1.4.).

#### 1.1. Computing cell to cell variability in gene expression with Shannon entropy

The Shannon entropy (H, measured in bits) is as a measure of how heterogeneous a probability distribution is in a given interval of measurements. Figure S1 illustrates how we can expect entropy to behave for different distributions.

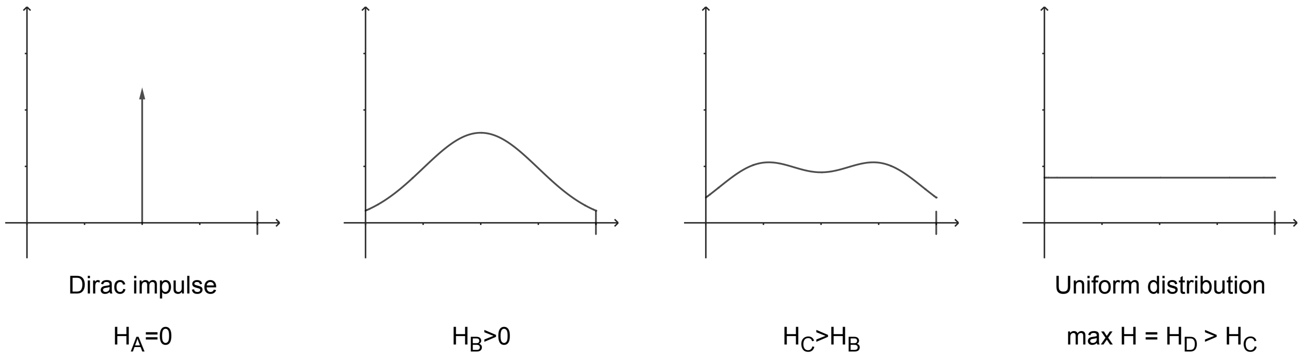

Figure S1: Illustration of Shannon entropy H for different distributions in the same interval. A Dirac impulse (A) has an entropy of 0, as all the probability is concentrated in a single point and there is no heterogeneity. A uniform distribution (D) is the most heterogeneous distribution possible within the range, and has the highest possible entropy for it. Between, these extremes, the more heterogeneous the distribution is, the higher the entropy.

This entropy was first defined over discrete distributions as $H(\chi) = -\sum_{x\in\chi} p(x)\log_2 p(x)$, where $\chi$ is the probability distribution for which the entropy is computed values and $p(x)$ is the probability of observing the value $x$ according to $\chi$, and any value for which $p(x) = 0$ is ignored in the sum.

While this definition can be generalised to continuous distributions, $-\int p(x)\log_2(p(x))dx$ usually converges to $\infty$ as a continuous distribution has infinite information in most cases. Therefore, the simplest way to measure the information of a continuous distribution is to discretise it, by dividing the range of possible values into bins and computing the entropy over the resulting discrete distribution.

The choice of these bins will affect the resulting measured entropy. In order to correctly represent the entropy of a distribution, the the bins should be smaller than the features of the distribution (i.e. it needs to be smaller than the peaks and troughs of the distribution for information on these peaks and troughs to be measured). However, there is no "convergence" of the entropy, where choosing smaller and smaller bins would lead to virtually identical results: to the limit, reducing the number of bins leads to an integral and, as stated above, increase towards infinity. In practice, when measuring the entropy of a measured distribution, individual points are available and binned. Decreasing the size of the bins as much as possible would lead to having a single measurement per bin, in which case the entropy will be the maximum possible entropy for the number of measurements available, and that is not useful. Therefore, for continuous distributions, the bins are often arbitrarily chosen, and the entropy is comparative between different distributions with the same binning strategy. The binning strategy itself is a research topic on its own, but for this paper we used a set of constant bins equally spaced in log space to compare different distributions.

For flow cytometry measurements, more than 10,000 cells are available, and in our case, only 2 major phenotypes are expected. To capture the heterogeneity, we used 50 bins covering the whole range of measurement values for each case study, as illustrated in Figure S2.

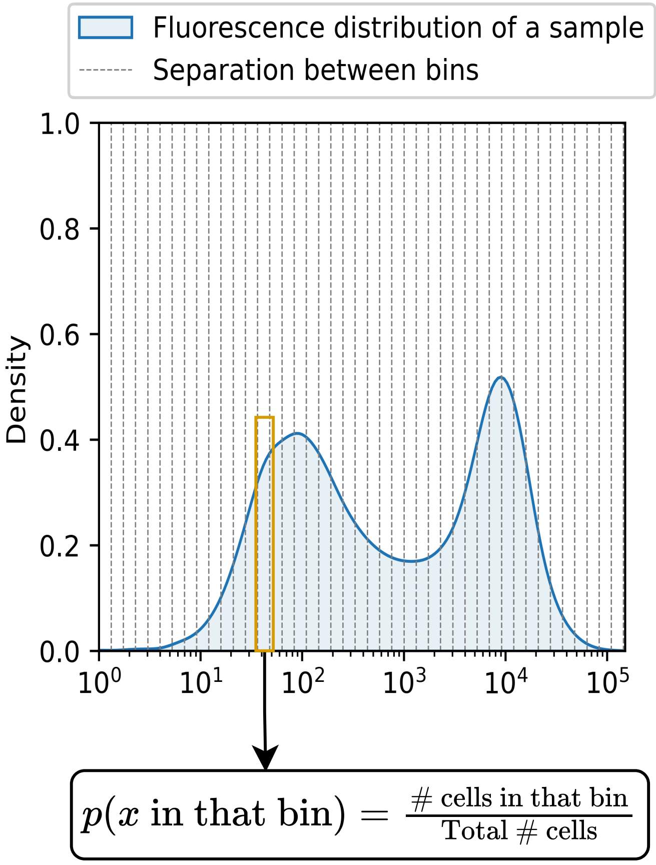

Figure S2: Example of the binning strategy used for the entropy measurements. The bins are equally spaced in log space, and cover the range of the measurements. The number of bins is much larger than the number of phenotypes expected (2), but much smaller than the number of cells measured (>10,000).

#### 1.2. Using entropy to compare the level of heterogeneity between populations

In addition to the definition of entropy, there is one important property of Shannon entropy that is relevant here: mixing two distributions increases their entropy. Intuitively, this can be understood as how mixing two distributions will make the result more heterogeneous. Mathematically, this is expressed as:

$$H(k \cdot \chi_1 + (1-k) \cdot \chi_2) \geq k \cdot H(\chi_1) + (1-k) \cdot H(\chi_2) \tag{S.1}$$

where $k$ is a number between 0 and 1, $\chi_1$ and $\chi_2$ are two distributions, and the left-hand side is the entropy of the mixture of the two distributions.

In particular, when the ranges of $x \in \chi_1$ and $x \in \chi_2$ do not overlap, the entropy of the mixture can be directly calculated as:

$$H(k \cdot \chi_1 + (1-k) \cdot \chi_2) = k \cdot H(\chi_1) + (1-k) \cdot H(\chi_2) - k \cdot \log_2(k) - (1-k) \cdot \log_2(1-k) \tag{S.2}$$

where $k$ being smaller than 1 means that this is indeed always larger than the mean entropy of the two individual distributions.

With this property, it is possible to determine what would happen to the entropy of a mix of two different cell populations, as depicted in Figure S3. On that figure, the x-axis represents the proportion of each population in the mixture, with 0 being 100% of the first population and 1 being 100% of the second one. The y-axis shows the entropy. Because of the aforementioned property, the lower limit of the entropy is the line joining the entropies of the two populations, and because there is a maximum entropy (the uniform distribution), any mix of two distributions must have an entropy somewhere within the depicted quadrilateral. Further, if the two populations do not overlap, for example when considering two independent phenotypes, then the entropy of the mixture can be calculated as in equation S.2, and the resulting entropy is shown with the blue curve.

The distributions of interest for this paper are the distributions of fluorescence of cell populations. In particular, we are comparing the distributions obtained in continuous cultures to individual phenotypes. These phenotypes are measured either as the autofluorescence or fluorescence of the cells and have a baseline entropy. When contrasting them with a high entropy distribution from a continuous culture, we are measuring how much more heterogeneous the continuous culture is by comparison to the individual phenotypes. This measurement can be applied on multimodal distributions, is independent of the amplitude of the signal, and can be applied to multi-dimensional data if required. For these reasons, it can be more useful than standard statistics, such as the standard deviation or Fano factor, to determine the heterogeneity of a sample.

#### 1.3. Differences in fitness: the root of the Fitness-Entropy compensation effect

Let us now investigate the Fitness-Entropy compensation effect. To do so, we can start with a simplified case in a chemostat, where the growth rate of the cells is the only factor determining their fitness. To showcase the effect, it is sufficient to consider two cases:

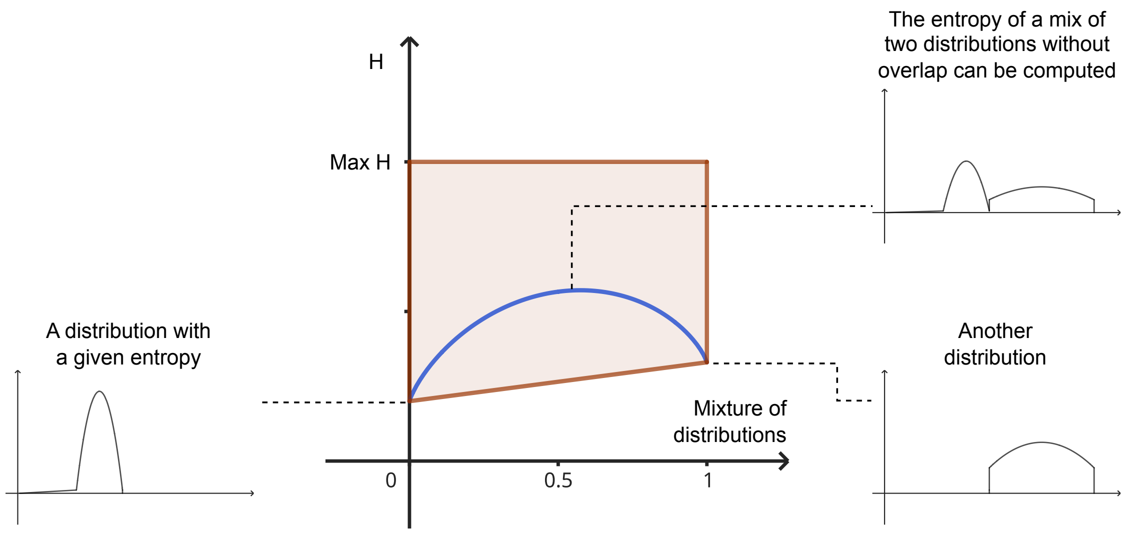

Figure S3: Entropy of the mixture of two different distributions; the x-axis represents the proportion of each population in the mixture, with 0 being 100% of the first population and 1 being 100% of the second one, and the y-axis shows the entropy. The quadrilateral on the entropy plot represents the area a mixture of any two distributions could have its entropy fall in. If those distributions do not overlap, like the ones depicted here, then the entropy of the mixture is the blue curve.

- A baseline case with two well-defined phenotypes, one induced and one uninduced, where all cells have the same growth rate.
- A case where we have the same phenotypes as previously, but the amount of induction affects the growth rate. Out of simplicity, we can consider that cells above a threshold expression level $T$ have a reduced growth rate $\mu_{lim}$, while cells below this threshold have the original growth rate $\mu_{max}$.

In the the baseline case, the two phenotypes $\varphi_1$ and $\varphi_2$ have the expected normal distribution as depicted in Figure S4.A and S4.B, respectively. By comparison, in the case of induction affecting growth, there is a selection effect: cells with a phenotype above the threshold $T$ have a reduced growth rate $\mu_{lim}$, and this decreases the relative abundance of the induced phenotype in the population, as shown in Figure S4.C and S4.D.

When looking at the uninduced phenotype, the distribution is nearly unaffected, as most cells are below the threshold $T$ and therefore have the original growth rate $\mu_{max}$. By contrast, the induced phenotype is selected against, and the selected distribution is a mix of the induced but unfit phenotype and a much larger amount of uninduced cells. These uninduced cells in Figure S4.D would normally be the tail end of the distribution in Figure S4, and are the noise of the distribution.

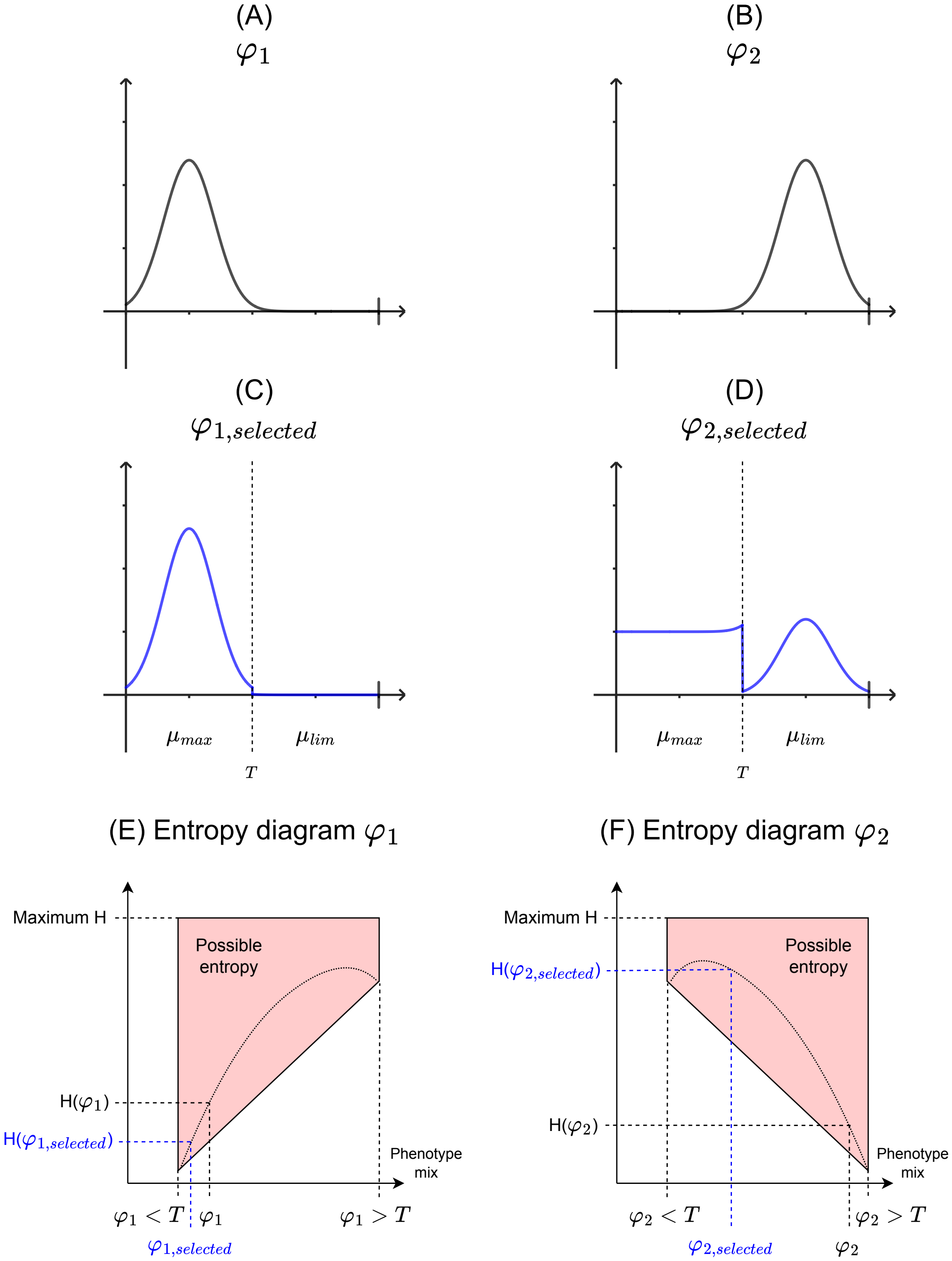

Figure S4: (A,B) Example probability distributions (not to scale) of fluorescence for two well-defined phenotypes, where the phenotype is not affecting fitness. (C,D) The same distributions, but where the phenotype is affecting fitness. Since they are well defined phenotype, we can expect most of the cells in $\varphi_1$ and $\varphi_2$ to be on the same side of the threshold $T$ separating the two phenotypes. The other side is mainly the tail end of the distribution, and is eliminated (C) or selected (D) as $\mu_{lim} < \mu_{max}$. (E,F) The entropy of the distributions compared to the entropies of the distribution split between the induced and uninduced regions. The entropy of the noisy part is much larger than the entropy of the main phenotype (by virtue of being noise).

Let us now look at the effect of this selection on the entropy of the distributions. The entropy of each distribution can be split into the entropy of the uninduced region and the entropy of the induced region, according to equation S.2, since we are considering no overlap. In practice, we can expect some overlap due to measurement errors and imperfect correlation between fluorescence and growth phenotype, but the principle holds. For each phenotype, the entropy of the region containing the bulk of the distribution ($H(\varphi_1 < T)$ and $H(\varphi_2 > T)$) will be quite small in comparison to their counterpart ($H(\varphi_1 > T)$ and $H(\varphi_2 < T)$), as the bulk of the distribution represents a defined phenotype, while the tails of the distribution are mainly noise and essentially flat, bringing their entropy close to the maximum possible entropy.

In Figure S4.E, it can be easily seen that selecting the main phenotype slightly reduces the entropy of the distribution as the portion of the distribution that is eliminated is the noisy one. However, because the noise for a well-defined phenotype is already a very small portion of the total distribution, this may not be measurable.

In Figure S4.F, we can see that, by contrast, selecting the noise instead of the induced phenotype naturally leads to an increase in entropy instead. This is the Fitness-Entropy Compensation mechanism and explain how entropy can be a measure of how much a phenotype affects fitness.

#### 1.4. Applying the Fitness-Entropy compensation effect to our case studies

There are two additional points to consider when determining whether the concept of Fitness-Entropy compensation should apply to a real system. First, in the simplified case of the previous section, only a single transition between two phenotypes was considered. In practice, the growth rate probably changes continuously with the phenotype. Second, we only considered selection of phenotypes, but there was no mention of cells switching from one phenotype to another.

##### 1.4.1. A fitness cost curve instead of a single transition between two phenotypes

In practice, we usually measured a continuous decrease in fitness with the measured phenotype, rather than two clear phenotypes with a clear threshold. To extend the previous reasoning, it is helpful to consider several discrete phenotypes rather than a smooth transition, knowing that if the principle holds for any number of phenotypes, then it holds to the limit if one considers the observed continuous decrease. Several discrete phenotypes could be depicted as in Figure S5.

To determine what is the entropy of such a distribution, equation S.2 can be generalised to any number of phenotypes:

$$\begin{aligned} H\left(\sum_{i=1}^{n} k_i \cdot \chi_i\right) &= \sum_{i=1}^{n} k_i \cdot H(\chi_i) - \sum_{i=1}^{n} k_i \cdot \log_2(k_i) \qquad \text{, where } \sum_{i=1}^{n} k_i = 1 \\ &= \text{mean}(H(\chi_i)) - \sum_{i=1}^{n} k_i \cdot \log_2(k_i) \end{aligned} \tag{S.3}$$

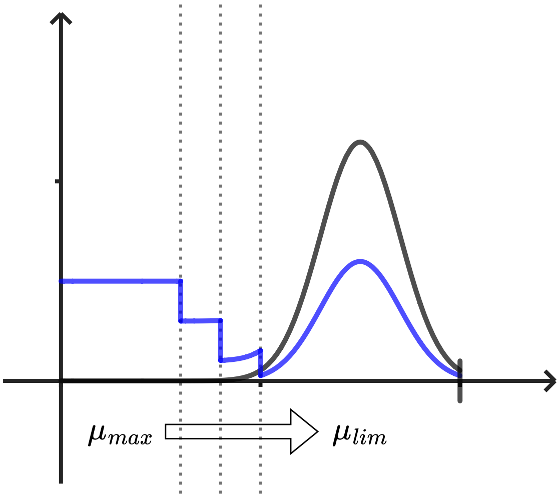

Figure S5: Entropy of a distribution with several phenotypes. In black the phenotype before selection. In blue, the phenotype after selection, where each successive region from left to right has a lower growth rate than the previous one.

This cannot be represented graphically as in Figure S4, but the same principle applies: the entropy of the combined distribution will be slightly higher than the mean entropy of the individual parts of the distribution. Since the case of interest is the one where the main phenotype is selected against and phenotypes in the tail of the distribution are amplified, the entropy of the combined distribution will always be higher than the entropy of the phenotype without selection.

##### 1.4.2. Switching between phenotypes

The second point to discuss is the switch of cells between the different phenotypes. Previously, the region selected against was always represented as having a smaller amount of cells. However, if cell lineages were to just switch once to a given phenotype then stop there, only the fastest growing phenotype would be present, over-growing any other. In practice, cells are traveling between phenotypes stochastically, and there should be a dynamic equilibrium between the faster growing phenotype that switches to the slower growing phenotype and the slower growing phenotype being washed out. A mathematical model of this equilibrium could be:

$$\frac{d\varphi_1}{dt} = \mu_1\varphi_1 - r_1\varphi_1 - D\varphi_1 + r_2\varphi_2 \qquad = 0 \text{ at steady state} \tag{S.4}$$

$$\frac{d\varphi_2}{dt} = \mu_{lim}\varphi_2 - r_2\varphi_2 - D\varphi_2 + r_1\varphi_1 \qquad = 0 \text{ at steady state} \tag{S.5}$$

where $\varphi_1$ and $\varphi_2$ are the concentrations of the two phenotypes, $D$ is the dilution rate, $\mu_1$ is the growth rate unlimited by the phenotype (lower than $\mu_{max}$ due to nutrient limitation), and $r_1$ and $r_2$ are the switching rates between the two phenotypes. If $r_2$ is nearly null, then $r_1 + D > \mu_{max}$ leads to the first phenotype being washed out, and the second follows, making the Fitness-Entropy compensation impossible when there is little noise and strong, fast induction. This explains why the Fitness-Entropy compensation effect was not observed for the toggle-switch, for example.

This model does not consider the environmental feedback that was observed conjointly with the Fitness-Entropy compensation effect, that would amplify any switching and ensure both populations can survive.

##### 1.4.3. A look at the numbers for the different test cases

We've seen that, in order to fully prove the fitness-entropy compensation effect, we need to show that the entropy of the selected phenotype is larger than both the entropies of the individual phenotypes. The entropies of the T7, ydcS, and glc3 systems were measured in different conditions to assess the validity of the effect more quantitatively. The results are presented in Figure S6.

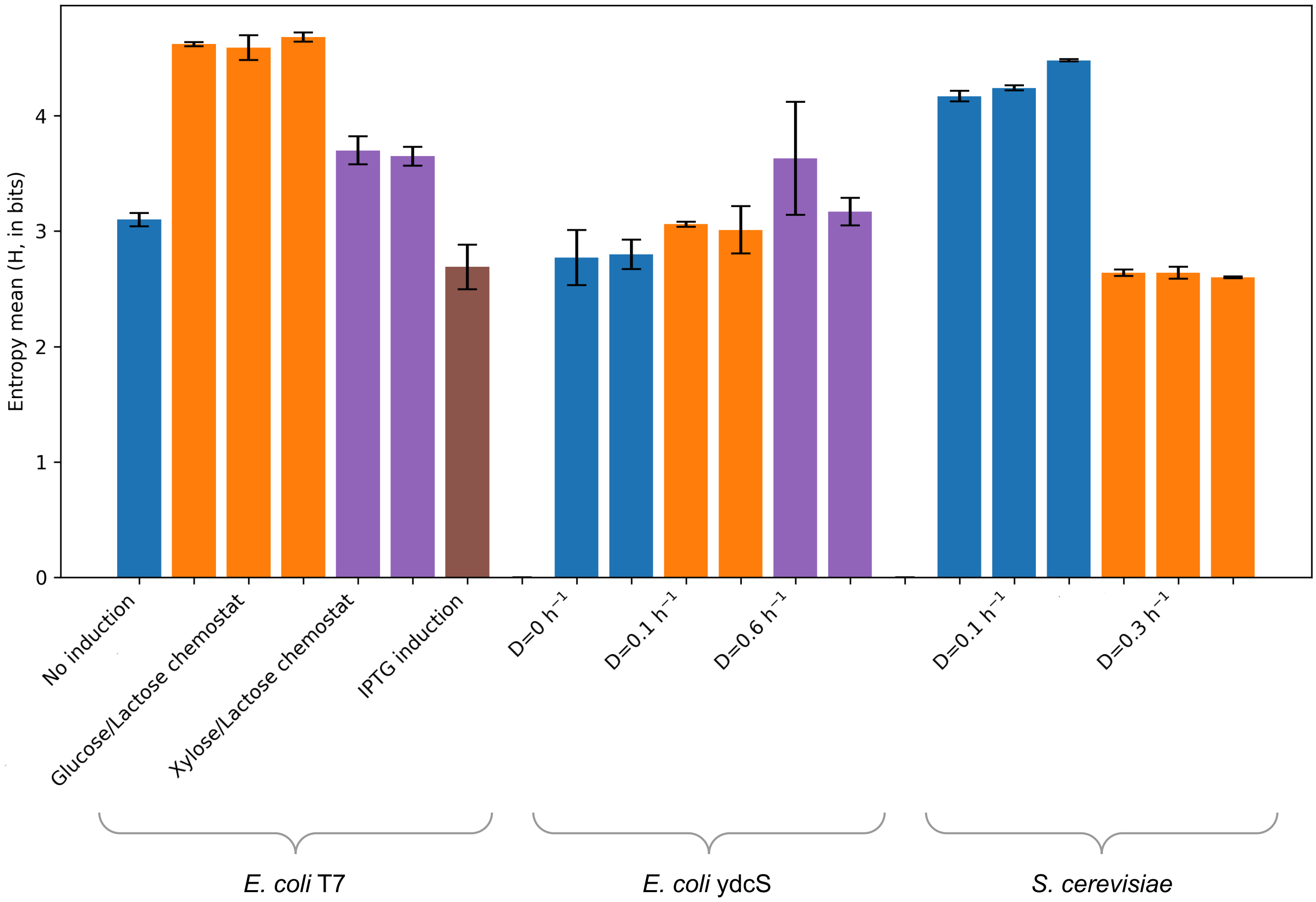

Figure S6: Entropy of the different systems measured in different conditions. Error bars are three standard deviations on 6 samples (technical replicates), except for the Uninduced and IPTG induced samples for the T7 system, which only have 2 samples each and come from flasks instead of bioreactors. Biological replicates are juxtaposed and in the same color. The entropies of the conditions where we would expect Fitness-Entropy are shown in the middle for each system. For the T7 system, the xylose/lactose and IPTG conditions are both purely induced phenotypes but have different entropies, primarily because the noise (measured as uninduced) in the xylose/lactose condition becomes more prevalent over time as cells are washed out of the bioreactor. For the ydcS system, D=0 $h^-1$ indicates the stationary phase in bioreactors without any feeding. There is no Fitness-Entropy compensation for that system, and the entropy at 0.1 $h^{-1}$ is in between the other conditions. For the yeast, only one of the unstressed reference point is available (0.3 $h^{-1}$), but it is much lower than the entropy observed in chemostat at a lower dilution rate (0.1 $h^{-1}$).

The entropy of the pure phenotypes for the T7 system was measured as T7 without inducer on one side, and as T7 with IPTG (in flask) or with xylose+lactose feed (from our bioreactors) on the other side. By comparison, the entropy of the chemostat on glucose+lactose, where we claim Fitness-Entropy compensation, is 25 to 50% higher than the entropy of the pure phenotypes[1], showing that we are indeed measuring the Fitness-Entropy compensation effect.

For the ydcS system, there should be no noticeable Fitness-Entropy compensation effect, as the stressed phenotypes have a higher growth rate than the dilution rate and no significant trade-off was observed. As expected, the entropy of the low dilution rate (D=0.1 $h^{-1}$) is in between and of a similar magnitude to the entropies of the pure phenotypes (D=0 $h^{-1}$, a stationary phase with no feeding causing starvation stress, and D=0.6 $h^{-1}$, a chemostat with a high dilution rate eliminating stress).

Finally, for the yeast, we were unfortunately not able to measure a purely induced phenotype in bioreactor. Batch cultivation with prolonged stationary phase to induce stress expression caused a bimodal distribution unlike what we observe in microfluidics. In flask, cells experienced successively conditions of the Crabtree effect (ethanol production), respiro-fermentation conditions (respiration of glucose and re-consumption of ethanol) and finally conditions with very low concentration of glucose. We are therefore not confident that this distribution was a reliable indicator of the population as it would happen if the glc3 system was induced by a uniform glucose starvation. The microfluidics measurements in stressed state are more representative and do indicate a well-defined phenotype, but we currently have no way to compare the entropy measurements from flow cytometry with microfluidics. As a result, we cannot conclude with full certainty that the Fitness-Entropy compensation effect is present in the glc3 system, but we believe that the increase in entropy between the high and low dilution rates (about 60%), the entropy at D=0.1 being higher than any entropy of a pure phenotype we measured, the uniform induction of the system in microfluidics, and the growth trade-off measured, are strong indicators in favour of the effect being present.

### Supplementary Note 2. Calibration of the automated flow cytometry

Three instruments were used for the automated flow cytometry measurements, two Accuri C6+ and one Accuri C6. For the T7 system in *E. coli* and the *glc3* system in *S. cerevisiae*, all three machines were used for different experiments, while for all other systems, only one machine per system was used. Since measurements are not identical between machines, we performed a calibration between them and transformed all the measurements of any machines into what one of our C6+ cytometers, which we are using as reference, would have measured.

To do so, identical samples of both yeast and bacteria were passed through either both C6+ machines, or our reference C6+ cytometer and the C6. For *S. cerevisiae*, the calibrations were made using pre-cultures of our GFP-producing strain with different levels of stress, due to starvation (cells in exponential growth or in stationary phase) or heat shock (culture in stationnary phase let 30min, 1h and 1h20 at

[1]Note that the difference with the xylose is only so low because the background noise of the cytometer (up to 50 events/µL) is larger in that case (only 300 events/µL because of the washout of the population) than in other bioreactors (about 1000 events/µL). This increases entropy as the noise does not overlap with the measurements of GFP.

37°C). For the *E. coli* T7 system, the calibrations were made using cultures on defined medium with glucose, with or without induction of the T7 system with IPTG.

The measurements were split in percentiles and a correlation was determined between the percentiles of the reference and the other cytometer, for both forward scatter (FSC-A) and fluorescence (FL1-A), which are the measurements we use for our analyses. The codes are available in the files `ConversionFromC6.py` and `ConversionFromC6plus_Left.py`.

For the Accuri C6 to C6+ transformation, the correlation that best fitted all data was a power law of the form $y = ax^b$, where $y$ is the measurement of the reference cytometer, $x$ is the measurement of the other cytometer, and $a$ and $b$ are parameters fitted to the data. A linear fit would have suffised for the FSC-A measurements and for FL1-A for *S. cerevisiae* (Figure S7.A), but for the T7 system the relationship was not linear at all (Figure S7.B). We chose our fit so that high fluorescence values were correct, while low values in the C6, gave an over-estimated fluorescence after transformation. The high values were prioritized because they are the ones used to determine the growth rate in segregostat. Nevertheless, the exponent in this power law means that the determination of growth rate for the T7 system needs to be adjusted to account for the non-linearity of the correlation.

A. C6 to C6+ Yeast Fluorescence　　B. C6 to C6+ *E.coli* T7 Fluorescence

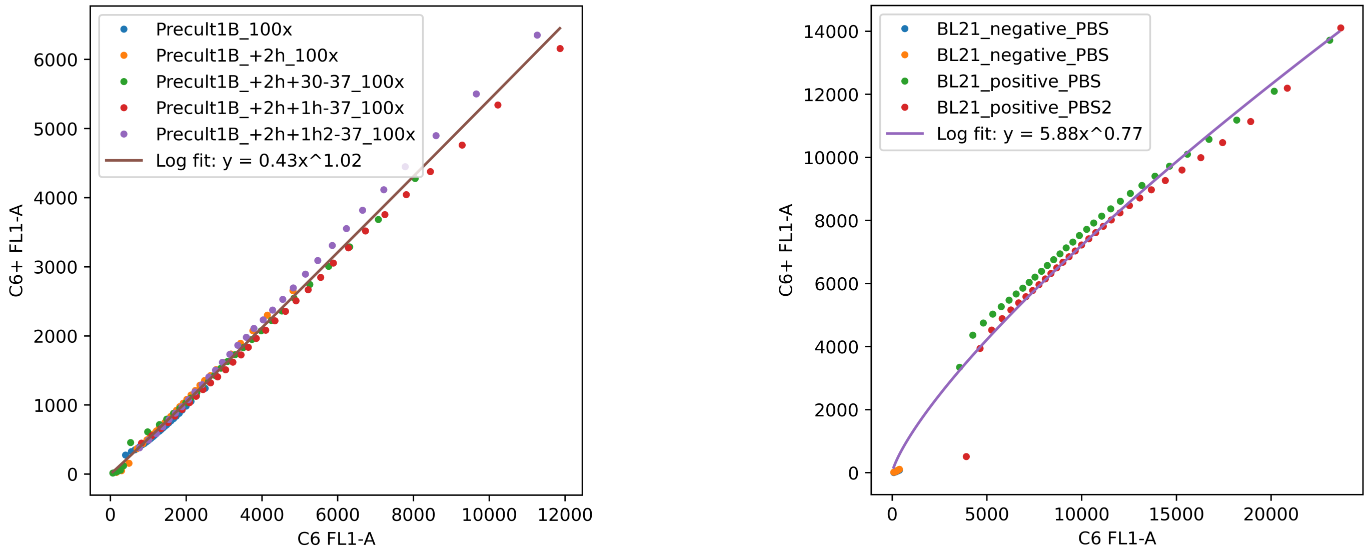

Figure S7: Calibration of the Accuri C6 cytometer with the Accuri C6+ cytometer. The data points are the percentiles of the measurements, and the curves are the fitted correlations.

For the transformation between the two C6+ machines, we used a linear correlation for all measurements, although the correlation for the T7 system fluorescence was not as good as others (Figure S8). This is likely because, for this system, the cells are very close to the detection limit of the cytometer, and different amounts of noise are measured by the two machines. This doesn't seem to affect the entropy significantly, however, since the entropy for each replicate of the Xylose/lactose and Glucose/lactose experiments were very similar despite coming from different machines (Figure S6).

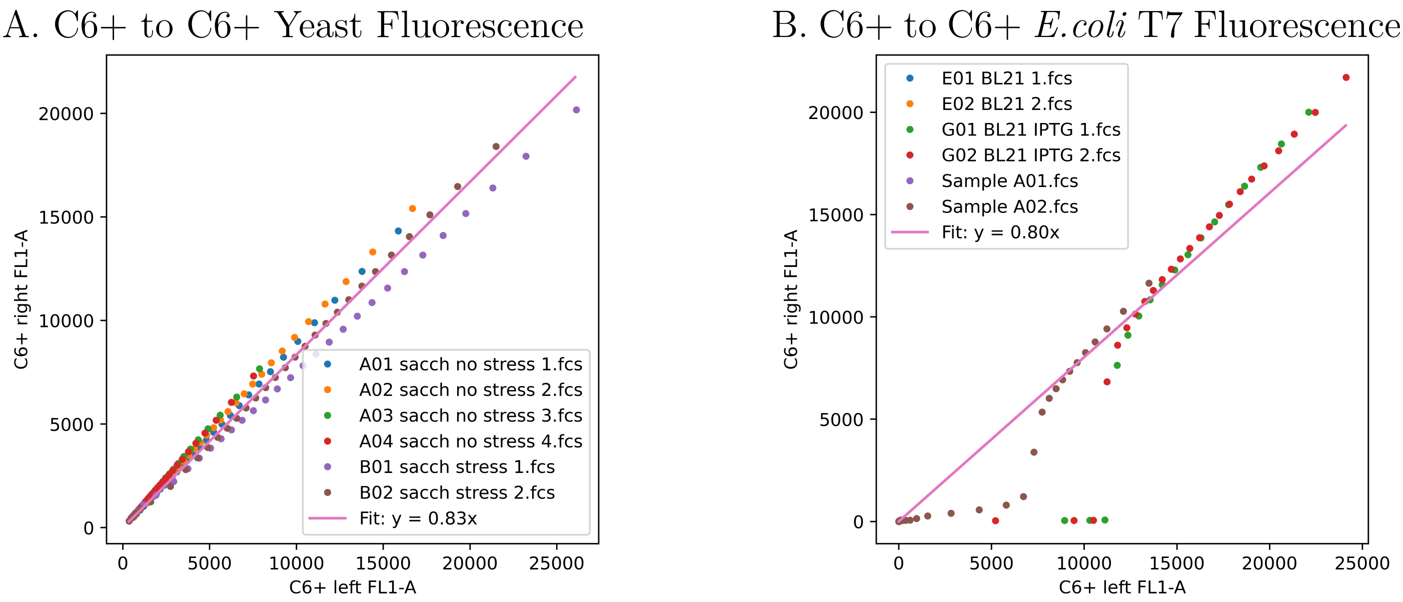

Figure S8: Calibration of the Accuri C6+ cytometer with another Accuri C6+ cytometer. The data points are the percentiles of the measurements, and the curves are the fitted correlations.

### Supplementary Note 3. Determination of the trade-off curves based on Segregostat experiments

#### 3.1. Basic principles of the method

To determine fitness, we needed to determine the growth rate as a function of the measured phenotype. To do so, we made use of Segregostat measurements, where induction is periodic instead of constant. Thanks to the periodicity, after each induction is a relaxation phase where no fluorescence is produced (see Figure S9.A). We used this relaxation phase to estimate phenotype growth rates; this estimation relies on a few assumptions:

- The growth rate is constant during the portion of relaxation phase considered. Technically, the cells are changing phenotype (recovering from stress, for example), but we used data from only up to 1.5 h to have 5 measurements and decrease the influence of the change in phenotype.
- During relaxation, no fluorescence is produced, and the main source of decrease is dilution. In other words, the decrease in fluorescence is only due to growth, and calculating it is equivalent to calculating the growth rate. This is pertinent because protein degradation is negligible compared to growth in most cases [2].
- The function of growth rate with respect to fluorescence is monotonously decreasing. What this means effectively is that, when two phenotypes are measured at time $t$ and $t + \Delta t$, a cell with a lower fluorescence than another at time $t$ will also have a lower fluorescence at time $t+\Delta t$ because it is growing faster, hence diluting the fluorescence more, and there is no intersection between the trajectories of the cells. This assumption underlies the entire strategy we are using and we will

come back to its validity. When there is a trade-off between growth and gene expression, this is a reasonable assumption as a higher fluorescence means more protein production, hence less growth.

Using these assumptions, we can measure the growth rate of a population by following the fluorescence of different portions of the population over time: if the the upper quartile of the population halves its fluorescence in 4 hours and the lower quartile halves it in 2, then the growth rate of the lower quartile is 0.35 $h^{-1}$, and twice that of the upper quartile (Figure S9.B).

In practice, the population was split in the 90 percentiles between 5 and 95% of the fluorescence distribution in order to capture the full range of growth rates. The fluorescence was followed over five successive automated flow cytometry measurements, that is roughly an hour, to avoid too much phenotypic change over the measurement period. This was done for multiple relaxation phases and for multiple biological replicates, to determine a Hill function (representing the inhibition of growth by fluorescence) that would correctly approximate the growth rate as a function of fluorescence (Figure S9.C for a single biological replicate). This trade-off curve can then be used to determine the fitness of a population based on its fluorescence measurements.

The code for this procedure is available in the file `Approx_mu_fFLUO_Seg_functions.py` (`decay_percentiles` function), it provides an estimate for the growth rate as a function of fluorescence at the start of the time interval considered for each percentile. It is used in the file `Approx_mu_fFLUO_Seg_fitfct.py` to find a function fit. This code takes into consideration the methods to mitigate various errors as described below.

#### 3.2. Mitigation of various sources of error

While this basic method is useful, there are several sources of error that can impact the measurements significantly. Some of them were accounted for in our data treatment to improve the accuracy of the results, while some minor factors cannot be corrected for accurately.

**Bigger cells have a higher fluorescence,** but do not necessarily grow slower because they also have more resources available. Instead, the measurement of interest is the fluorescence concentration in each cells. Therefore, instead of using the raw fluorescence, we used the fluorescence divided by the forward scatter (FSC) measurement, which is a proxy for the size of the cells. For yeast, FSC is linear with respect to cell volume [3], and this correction is valid. For bacteria, the relationship between FSC and cell volume is harder to establish, but at the very least, very stressed cells that filament and have a much higher FSC do not inflate the fluorescence excessively if this normalisation is applied.

**The auto-fluorescence of the cells** implies that fluorescence measurements do not follow a simple decaying exponential (Figure S10). Rather, they follow a decaying exponential shifted up towards the auto-fluorescence ($Fluo = Fluo_0 \cdot e^{-\mu t} + Fluo_{auto}$). Without taking that into account, the growth rate would be systematically underestimated. To correct for this, we subtracted the mean auto-fluorescence of a blank sample without induction from the fluorescence measurements. Moreover, the auto-fluorescence is not the same for all cells. To avoid confusing fluorescence decay with auto-fluorescence, we excluded any fluorescence point that entered the range of auto-fluorescence from our measurements.

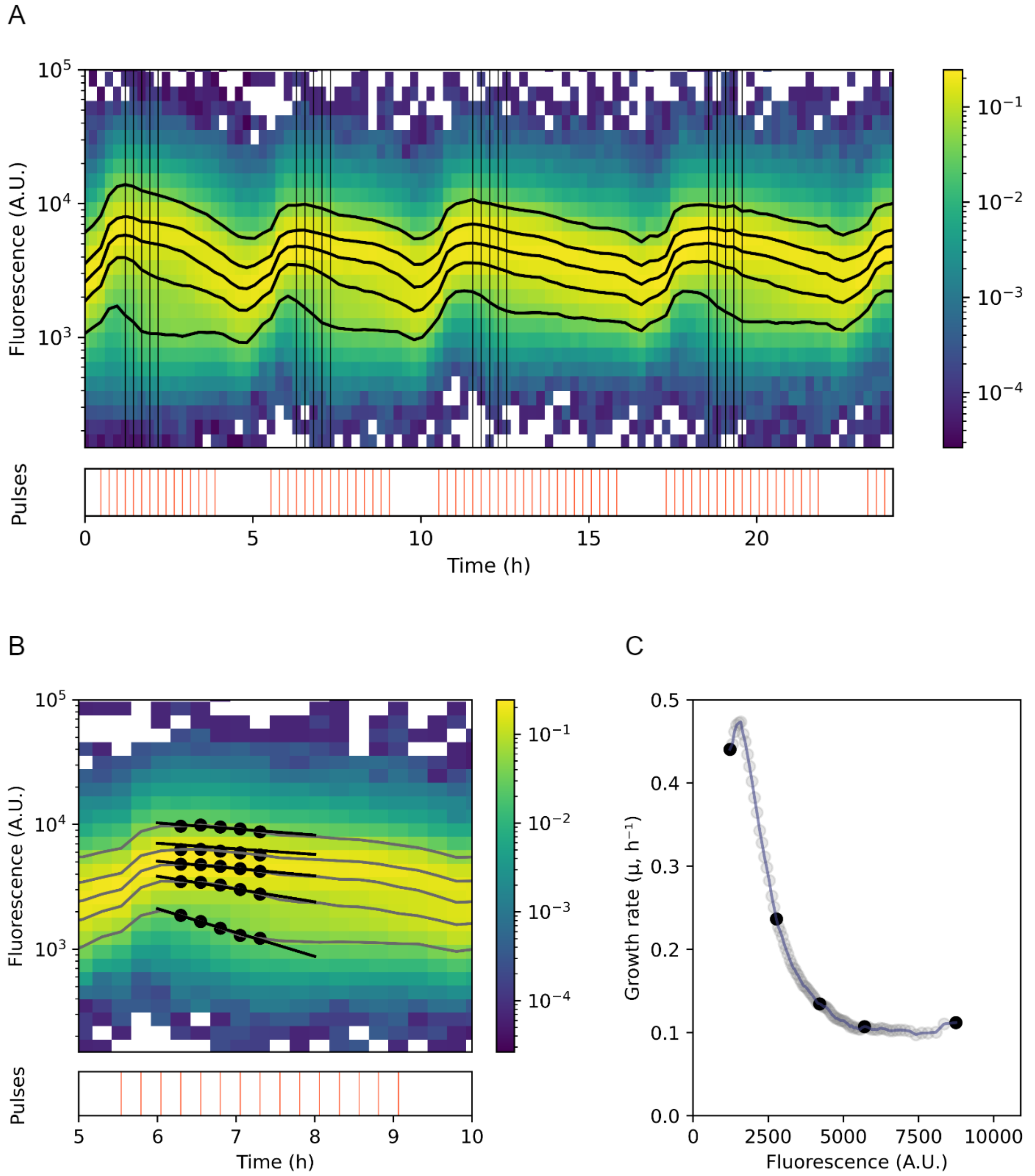

Figure S9: A. The time scatter of a segregostat experiment for the yeast, showing the measured fluorescence over time, lighter colors indicate more cells at that time and fluorescence. The lines in the 'pulses' rectangle indicate when glucose was pulsed to reduce the stress levels. The black vertical lines on the scatter plot indicate the ample times used to compute the trade-off curve. The black lines following the population indicate percentiles 5, 25, 50, 75 and 95. Periodic induction of stress and relaxation phases are produced. B. For each relaxation phase, exponential curves (black lines) were fitted to the five selected time points (black dots, along the same percentiles (gray lines) shown in (A)). Note that these fits do not account for the mitigation of various sources of error described in the next section. C. Trade-off between gene expression (fluorescence) and growth rate, as derived from the fitted exponential curves. Each round dot corresponds to a fluorescence percentile; black dots indicate the results for the specific percentiles illustrated in panels (A) and (B).

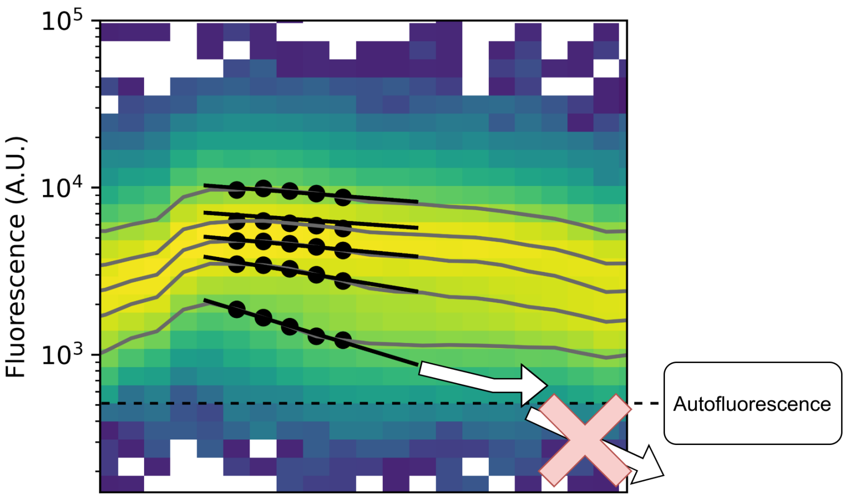

Figure S10: When fitting a simple decaying exponential, we assume the fluorescence is decaying towards zero. However, the fluorescence measurement is a sum of fluorescence due to GFP and auto-fluorescence. The decaying exponential should be going towards the autofluorescence (roughly 500 for yeast), so that going from 2000 to 1000 in fluorescence represents a 70% decrease in the fluorescence measurement due to GFP, instead of only 50%.

**Because all portions of the populations do not grow at the same rate,** the abundance of each section of the population (in our case, each percentile at a given time) changes over time: the slower, high fluorescence cells are washed-out while the faster, low fluorescence cells grow and take over the reactor (Figure S11). This means, for example, that the 25th fluorescence percentile at a given time may correspond to 30% of the population one hour later instead of 25%. Therefore, with the basic method, we would over-estimate the growth rate of each sub-population. This was taken into account by iteratively estimating the growth rate of the population, then using it to measure which percentiles at a given time correspond to which percentiles in the other samples, then re-computing the growth rate based on these new percentiles until the change in growth rate was negligible.

We investigated the effect of this correction for the *glc3* system in *S. cerevisiae* and found that it changed the estimated growth rate by up to 20%, showing that it is an essential step to obtain a good measurement.

Notably, in order to compute the change in abundance of each percentile of the population in the bioreactor, the dilution rate of the reactor is used. In our setup, the volume of the bioreactor may change by $\pm 5\%$ between experiments, and the tube used for feeding being autoclaved and wearing out may add error on top of that. To evaluate the impact of this error, we performed the calculations for the second replicate of the *glc3* system, which has the most relaxation phases, with dilution rates of 0.1, 0.125 (the actual dilution rate), and 0.15 $h^{-1}$. The resulting Hill function parameters changed by less than $10^{-5}\%$, showing that this error is negligible.

**The assumption that the growth rate is monotonously decreasing with fluorescence** is reasonable, but not always measurable. In the range of fluorescence where the growth rate clearly decreases, everything works as intended. However, for low fluorescence values, the growth rate should roughly constant, and an increase in growth rate is often measured instead (as can be observed in Figure S9 for example). Similarly, for the highest fluorescence values, the growth rate reaches a minimum and, in some cases, we measure a small portion of the population with a

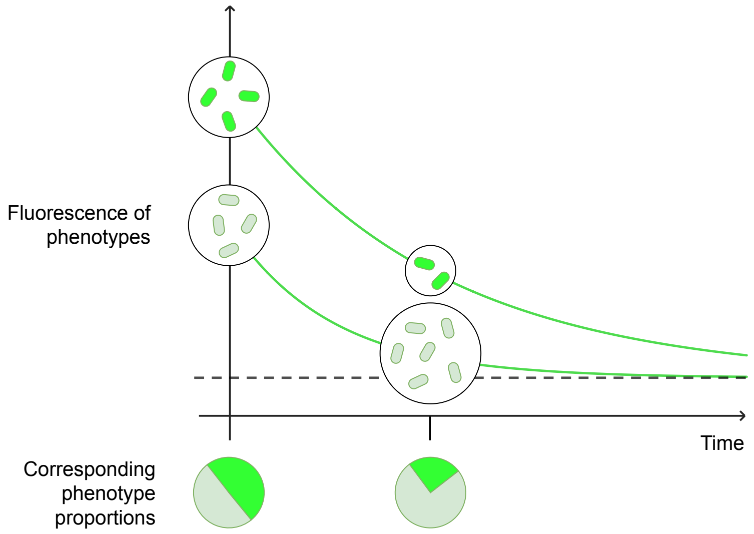

Figure S11: As the fluorescence decreases, faster-growing phenotypes also increase in proportion in the population. Based on the estimated growth rates, this change in proportion is estimated, and taken into account to correct the growth rates.

small increase in growth rate. When correcting the percentiles, we replaced these aberrant values with the maxima or minima of the measured growth rates, and we did not use these points to fit the Hill function for the trade-off curve.

**The growth rate of the cells may change over time,** as we are measuring it during the recovery from stress. To limit this effect, we only used about an hour of measurements, as early as possible when the fluorescence starts to decrease. The cells may be recovering anyways and it is not possible to accurately estimate this effect, we simply know that we may be over-estimating the growth rate of the stressed cells because they are recovering.

**Our cytometers are not linear with respect to fluorescence for bacteria,** as mentioned in section Supplementary Note 2.. For yeast, this does not seem to be a problem, but for the T7 system in *E. coli* BL21, discrepancies were observed. The same discrepancies were not observed when measuring the *ydcS* system in both C6+ instead, suggesting the problem is specific to BL21.

The relationship between our C6 and C6+ cytometers in particular is interesting as the correlation is of the shape $y = ax^b$, where $y$ is the measurement of the C6+ cytometer, $x$ is the measurement of the C6 cytometer, and $a$ and $b$ are parameters fitted to the data. Assuming that the relationship between actual fluorescence and any cytometer measurement follows a similar relationship, all measured growth rates would have an identical error factor of $b$. This was taken into account by measuring the average measured growth rate during the relaxation phase, and normalising the fitness-cost curve by the factor 'average growth rate/dilution rate'. We applied this correction for all bacterial systems. For the *ydcS* and *araB* systems, the resulting normalisation factors were 1.1 and 1.01, respectively, showing that the issue is quite small. For BL21, the normalisation factor was 1.7, showing large non-linearities between the reality and cytometer measurements.

**The T7 system in *E.coli* BL21 stops being induced at different times based on fluorescence.** Unlike for other systems, where the onset of recovery from induction was uniform accross the

population, for the T7 system, the lower fluorescence levels decreased in fluorescence while the higher levels were still increasing. Selecting samples too early therefore caused the apparition of negative growth rates foor high fluorescence values in our measurements, due to the fluorescence increasing. We set these values to 0 $h^{-1}$ when correcting the percentiles and did not use them to fit the Hill function, but if this part of the curve was negative, then the rest of it was underestimated. Similarly, when taking the samples too late, a high proportion (often 30 to 50%) of the population was already very low in fluorescence and couldn't be measured, and the rest of the population was likely already recovering from the stress, so taking that estimate would overestimate the growth rate.
To get a reasonable estimate, for each relaxation phase, multiple successive sets of samples were used (Figure S12). If the first are underestimated and the last are overestimated, then fitting the Hill function on all the points provides the best possible estimate. The adapted procedure to find a function fit is available in the file `Approx_mu_fFLUO_Seg_Ecoli_T7.py`.

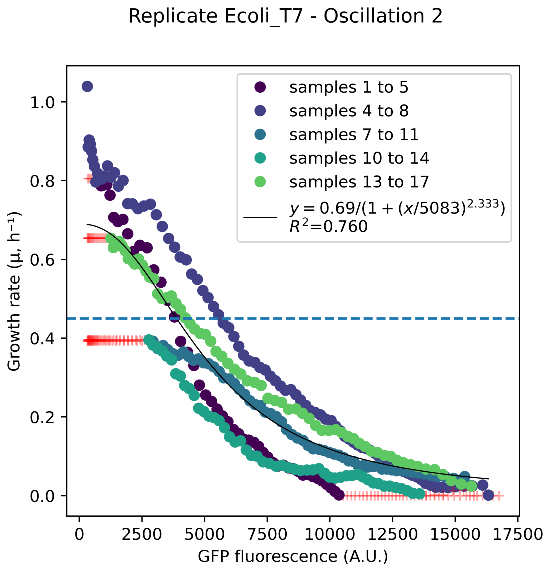

Figure S12: One relaxation phase of the *T7* system in *E. coli* BL21. Each color corresponds to one set of samples. The fitted Hill function was fitted on not all the relaxation phases of the replicate, though only one is shown here. The earlier measurements are under-estimated, the later ones are over-estimated, and the approximation is in between. Red '+' points were invalid and not used for fitting the Hill function. The dilution rate of 0.45 $h^{-1}$ used in the Segregostat is shown in blue dashes.

#### 3.3. Estimation of the trade-off curves for the four systems considered

##### 3.3.1. The *ydcS* and *araB* systems in *E. coli*

The cases of *ydcS* and *ara* in *E. coli* are examples where the trade-off between growth and fluorescence should be small or inexistent. The results are shown in Figure S13. Interestingly, for these two cases, correcting the final hill parameters by the dilution rate did not change the results much.

For the *ydcS* system, the measurements in chemostat show that the fluorescence can be much higher than what is obtained at 0.325 $h^{-1}$ when the dilution rate is lower (see Figure 6.A in the main paper).

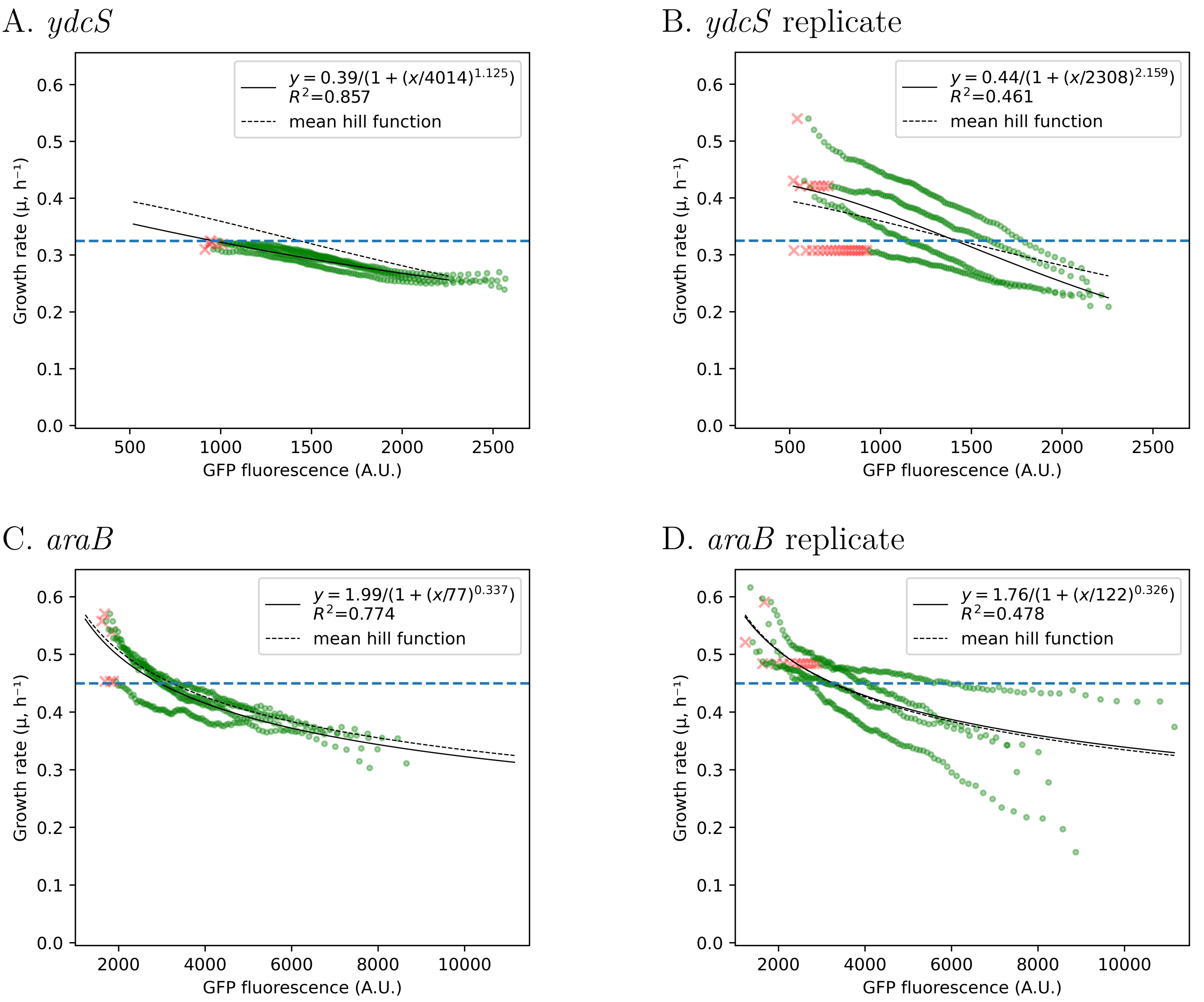

Figure S13: Growth rates measured for the *ydcS* and *araB* systems in *E. coli*. Each green point is a valid measurement of the growth rate as a function of fluorescence. No point was measured as <0 and replaced by 0 for the correction of percentiles (subpopulation proportions) based on the growth rate. Each red 'x' was before the maximum of the measured points, and none were after the minimum. This was inconsistent with the requirement that the curve be monotonic. Therefore, they were replaced by the maximum of the measured points for the correction of percentiles (subpopulation proportions) based on the growth rate. For the fitting, of the Hill function (full line), only the green points shown were used. The hill parameters were fitted independently for each replicated then averaged (dashed line). The horizontal dotted line is the dilution rates of 0.325 $h^{-1}$ for *ydcS* and $0.45^{-1}$ for *araB* used in the Segregostat. The measured average growth rates before scaling, based on the individual Hill functions, for each replicate, were 0.30 and 0.35 $h^{-1}$ for *ydcS* and 0.44 and $0.46^{-1}$ for *araB*. The scaling factor mentioned in the previous section is already applied in these figures.

Therefore, given the homogeneity of the population, an approximation of the trade-off function was obtained using chemostat data acquired at various dilution rates (Figure S14). The adapted procedure is available in the file `Approx_mu_fFLUO_varD_fitfct.py`. Only one replicate of chemostat at D=0.3 $h^{-1}$ have been performed in Bionet. Nonetheless, the values at D=0.3 $h^{-1}$ are close to the ones observed in segregostat. Furthermore, cultivations at the 3 considered growth rates have also been performed in

Dasgip and provides similar results (Figure S19). In addition, microfluidics measurements show only a slight reduction in growth rate with fluorescence. This is confirmed here, as we measure a growth rate that never gets below even 0.2 $h^{-1}$.

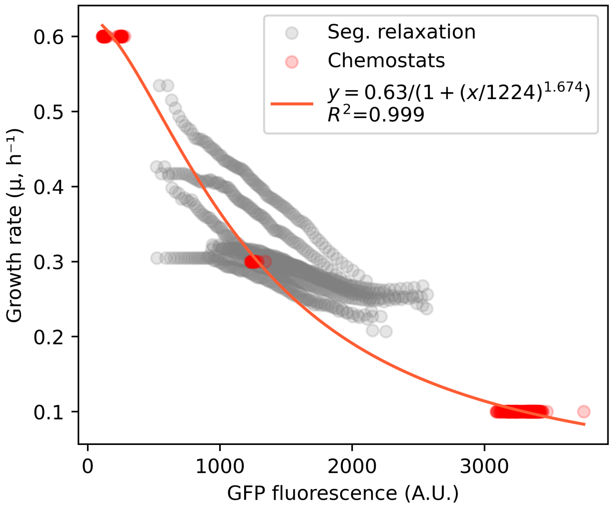

Figure S14: Trade-off function fitted on median fluorescence values obtained during chemostat cultivations at different dilution rate (see 'Approx_mu_fFLUO_varD_fitfct.py'). The points obtained with the method exploiting fluorescence relaxation in segregostat are indicated for comparison.

For the *araB* system, there should not be any trade-off between growth and fluorescence, as the maximum growth rate on arabinose only of the strain is higher than 0.5 $h^{-1}$ [4]. Similarly to *ydcS*, we show no growth rate below 0.3 $h^{-1}$.

For both systems, even though there should be no trade-off, we still measure a difference in growth rate with fluorescence, although it is much smaller than the cases with trade-offs. This can be explained by two factors:

- On the one hand, while there might be no trade-off, we are not measuring the maximum growth rate, but only the growth rate in the culture for given fluorescences. Even with maximum growth rates above the dilution rates, some phenotypes may still be unable to compete with others for limited nutrients, and therefore be slightly disadvantaged. Considering the populations observed are unimodal, however, this effect should be minor.
- On the other hand, if we are expecting no trade-off, then the unimodal, homogeneous populations measured have their variance mainly produced by noise in the measurements and noise in GFP expression. The closer we are to measuring only noise, the more the assumption that cell trajectories are not crossing each other is violated.

Considering these facts, since we know there should be no trade-off, we can consider the minimum growth rates obtained for these cases as the minima that need to be exceeded to ensure there is, actually, a trade-off between growth and gene expression. These minima are 30 to 40% lower than the dilution rate, and below that we can reasonably expect the growth curves calculated with this technique to reflect reality, as is the case for the *glc3* system in *S. cerevisiae* and the *T7* system in *E. coli* BL21, where we observe a reduction of at least 80% for the lowest growth rates.

##### 3.3.2. The *glc3* system in *S. cerevisiae* and the *T7* system in *E. coli* BL21

First, for the *glc3* system in *S. cerevisiae*, a high trade-off is expected based on microfluidics cultivation. This trade-off is confirmed by the measurements in the Segregostat, as shown for three biological replicates in Figure S15. The growth rate as a function of fluorescence is well fitted by a Hill function, however the first replicate is slightly different compared to the others. Since it is the only one made with the C6+ cytometer, which was the model used for the chemostat experiments, we use the parameters of the first replicate as the trade-off curve in the main paper.

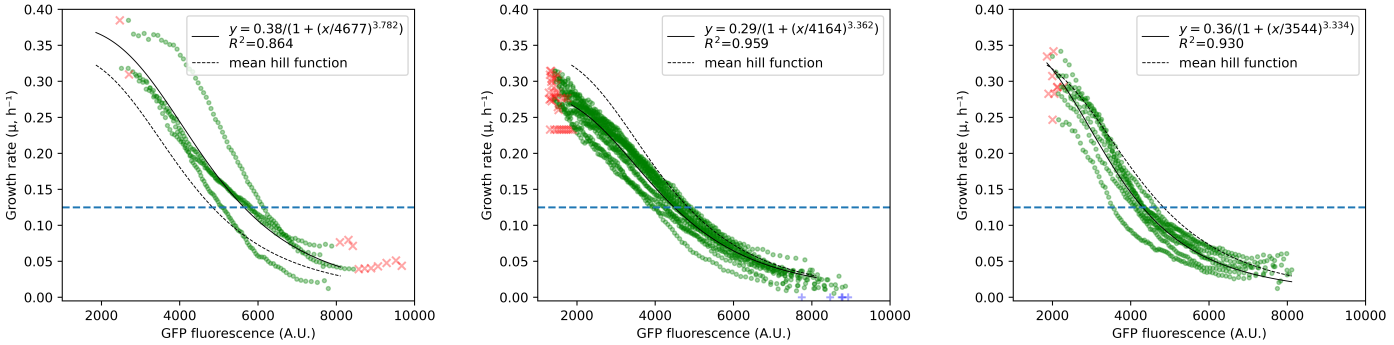

Figure S15: Fitness determination results for the *glc3* system in *S. cerevisiae*. Each green point is a valid measurement of the growth rate as a function of fluorescence. Each blue '+' was measured as <0 and replaced by 0 for the correction of percentiles (subpopulation proportions) based on the growth rate. Each red 'x' was either before the maximum of the measured points or after the minimum (and hence did not make sense since the curve needs to be monotonously). They were replaced by the maximum or minimum of the measured points, respectively, for the correction of percentiles (subpopulation proportions) based on the growth rate. For the fitting, of the Hill function (full line), only the green points shown were used. The hill parameters were fitted independently for each replicated then averaged (dashed line). The horizontal dotted line is the dilution rate of 0.125 $h^{-1}$ used in the Segregostat, and the measured average growth rates based on the individual fitted lines were 0.18, 0.16 and 0.13 $h^{-1}$ for the three replicates, respectively, in order of appearance.

Based on the fitted Hill function, we computed the growth rate of these populations and measured slightly higher values than the dilution rate of 0.125 $h^{-1}$. This may have (and probably has) two main causes. It can indicate that we are slightly overestimating the growth rate overall. However, for the *glc3* system, the relaxation phase happens when the culture is fed with additional glucose, which allows the biomass to increase. Therefore, another possibility is that the growth rate was actually higher than the dilution rate.

The *T7* system in *E. coli* BL21 is the system for which the measurements are the most imprecise due to the various caveats mentioned in the previous section, and we had to use a different measurement technique (see file `Approx_mu_fFLUO_Seg_Ecoli_T7.py`). Nevertheless, we obtained a reasonable estimate of the trade-off curve, as shown in Figure S12. The second replicate has more variance in the measurements but shows a similar resulting Hill function of $0.78 \cdot ({4815^{1.8}}/{4815^{1.8}} + x^{1.8})$.

### Supplementary Note 4. Characterization of the genetic toggle switches with and without trade-off

Individual phenotypes of the toggle-switch with influence on growth (i.e., with auxotrophy to its plasmid) were characterized before bioreactor experiments. The cultures were performed in a Biolector (M2PLabs, Germany), a microfermentation system that enables online monitoring of biomass and fluorescence in microtiter plates, as described below in Extended materials and methods. The measurments showed that the growth rate of the growing phenotype was $0.531 \pm 0.014$ (std) $h^{-1}$, while the non-growing phenotype had a growth rate of $0.295 \pm 0.010$ (std) $h^{-1}$ (data available in the supplementary dataset and code available in the `ToggleSwitchBiolectorAnalysis.py` file). The latter measurement is much smaller than the dilution rate used in bioreactor, and therefore using IPTG should wash the culture out.

As described in the main text, continuous cultures of both toggle-switch strains were carried out in lab-scale bioreactors. While the cultivation of the strain with growth control is shown in the main paper (Figure 4), that of the negative control strain, which has no effect on growth, is presented below (Figure S16).

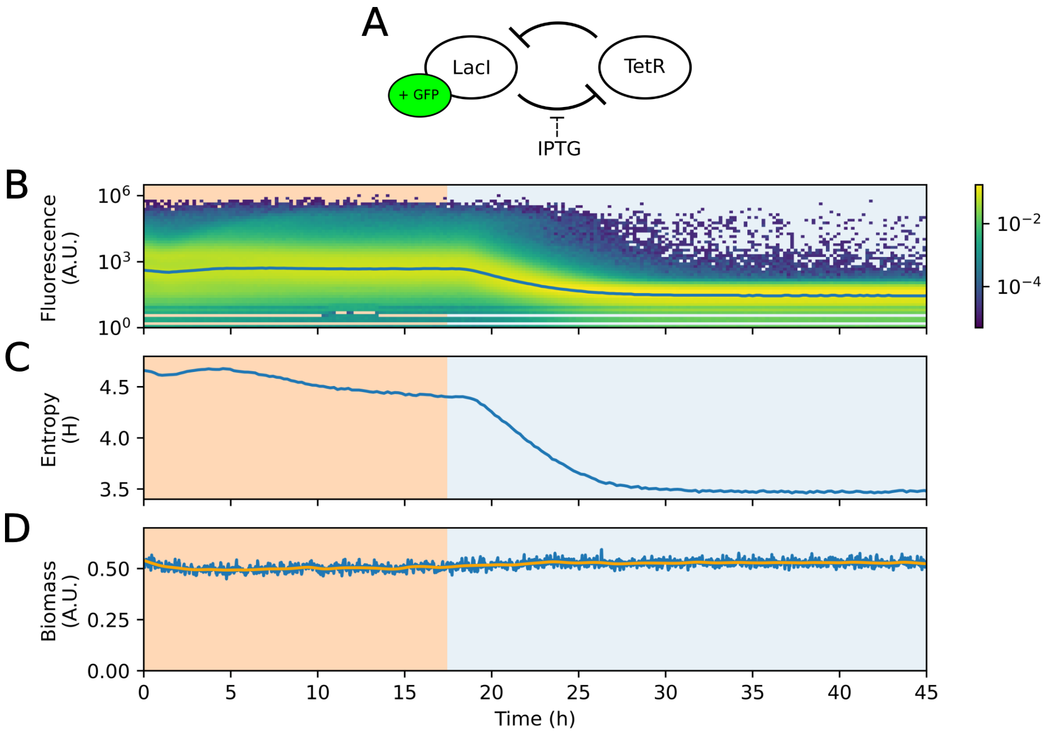

Figure S16: Continuous cultivation of the negative control toggle switch (without serine auxotrophy), upon addition of IPTG in the feed (blue background), the production of GFP was inhibited. (A) Schematic representation of the toggle switch. (B) Single-cell fluorescence distributions over time, the blue line indicates the median fluorescence. (C) Shannon entropy computed from the fluorescence distribution over time. (D) Biomass over time, the blue line indicates raw data and the orange line indicates smoothed data.

#### 4.1. Extended materials and methods

##### 4.1.1. Characterization in Biolector platform

Pre-cultures in defined medium from glycerol stocks were first made without inducer. These pre-cultures were then used to inoculate flasks with either 150 ng/mL anhydrotetracycline (less toxic than tetracycline) or 1 mM IPTG to produce pure cultures of each phenotype. These cultures were incubated overnight and washed twice in defined medium without sugar to remove the inducer. The OD was then adjusted to inoculate a Biolector plate at an OD of 0.5. In the biolector, each pure phenotype was grown in triplicate with and without inducer. For the growing phenotype, tetracycline was used instead of anhydrotetracycline in the Biolector at a concentration of 25 ng/mL, as it had been found enough to induce the phenotype but below the MIC. The non-growing phenotype was induced with 1 mM IPTG.

Growth curves for each well were produced by removing the baseline from the measurements then fitting an exponential to the exponential phase of growth by linear regression (see supplementary code).

### Supplementary Note 5. Selection of the general stress reporter for *E. coli*

We conducted a comprehensive search for suitable transcriptional reporters to detect the activation of the general stress response. Given that RpoS is the master regulator (transcription factor) of this response, we initially selected 26 genes from its regulon. These genes were chosen to represent the approximately 500 genes in the RpoS regulon based on the following criteria: (i) noise level [5], (ii) regulatory complexity (including $\sigma$S exclusivity, co-regulation by other transcription factors, position in the regulation network (regulator/regulated), (p)ppGpp dependency, etc.), (iii) sensitivity to RpoS concentrations [6], (iv) functions in response to various stress (pH, osmotic, etc.), (v) anti-correlation with growth, and (vi) correlation with antibiotic persistence.

The initial screening was performed using the Biolector platform to evaluate the basal expression levels, dynamic expression changes during the transition to stationary phase (i.e., entry into stressful conditions), and the dynamic range of expression (difference between basal and maximum expression levels) (Figure S17). Samples were taken during the exponential growth phase and late stationary phase for cytometry analysis and Shannon entropy computation (Figure S18). Based on these results, we shortlisted candidates using the following selection criteria: (i) expression levels (to be detectable by our flow cytometer), (ii) dynamic range (to clearly differentiate stress stages), and (iii) dynamic expression patterns during the transition to stationary phase. The selected reporters were further tested in a continuous cultivation device across various dilution rates, which correspond to different RpoS (i.e., stress) levels [7] (Figure S19). From these experiments, we ultimately identified YdcS as the optimal reporter. YdcS is a putative transport protein associated with natural and chemical transformations [8] and carbon flux distribution [9]. Its transcription is specifically initiated by $\sigma$S-containing RNA polymerase (E$\sigma$S) [8]. Follow-up analysis using Microfluidic Single-Cell Cultivation (MSCC) confirmed that YdcS is a reliable reporter of the general stress response (Supplementary Note 6.).

One may ask why we did not worked with RpoS transcriptional reporter directly. There are three mains reasons. First, the RpoS transcriptional reporter of the Zaslaver's collection seems deficient. Probably because the major rpoS promotor (rpoSp1) that is located within the nlpD gene, over 500 nucleotides upstream the rpoS gene, is missing in the plasmid [10–12]. Secondly, since few molecules of RpoS are produced per cell, detecting its transcriptional reporter with our cytometry setup would be challenging. Finally, RpoS transcriptional reporter may be a bit tricky since RpoS can accumulate without proper assembly into the active RNAP holoenzyme (E$\sigma$S) which directly regulates stress-related gene expression [11]. These challenges necessitated the use of an alternative approach. An ideal reporter gene would be exclusively regulated by E$\sigma$S and exhibit a simple regulatory system—criteria met by ydcS.

To verify that ydcS is a good indirect reporter of E$\sigma$S activity, we used a dual-reporter strain (*E. coli* MG1655 PrpoS::CyOFP1 PydcS::GFPmut2). This strain combines an RpoS transcriptional reporter integrated into the chromosome (capturing the major promoter) with a plasmid-based ydcS transcriptional reporter from the Zaslaver collection (see Materials and Methods in the main manuscript). Using microfluidics, we demonstrated that ydcS (GFP) expression increases under stressful conditions and correlates well with rpoS (CyOFP) levels (Figure S20). This finding confirms that ydcS is a reliable and convenient reporter for general stress response activation.

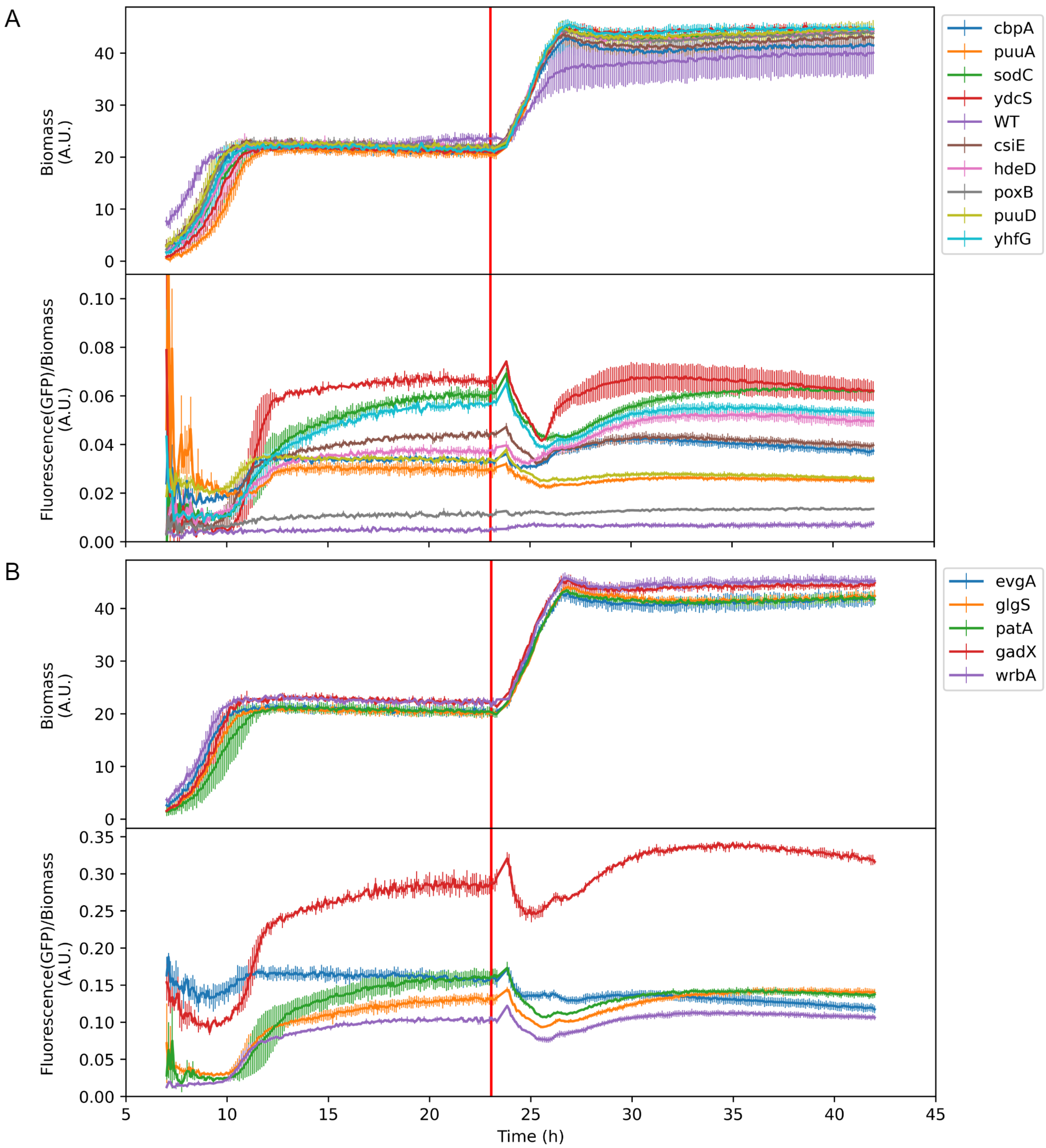

Figure S17: Example of visualization of data from the Biolector screening for reporters with low to intermediate expression levels (**A**) and reporters with higher expression levels (**B**). The top panel represents the biomass over time, while the second panel shows the relative fluorescence (GFP signal normalized by biomass) over time. Due to the low initial biomass, measurements are noisy at the beginning of the culture. Therefore, data collected before most wells display monotonal exponential growth have been excluded. Vertical lines represent the standard deviation (n = 3). The red vertical line at approximately 23 hours marks the time of sampling for cytometry analysis and the addition of glucose to the micro-wells.

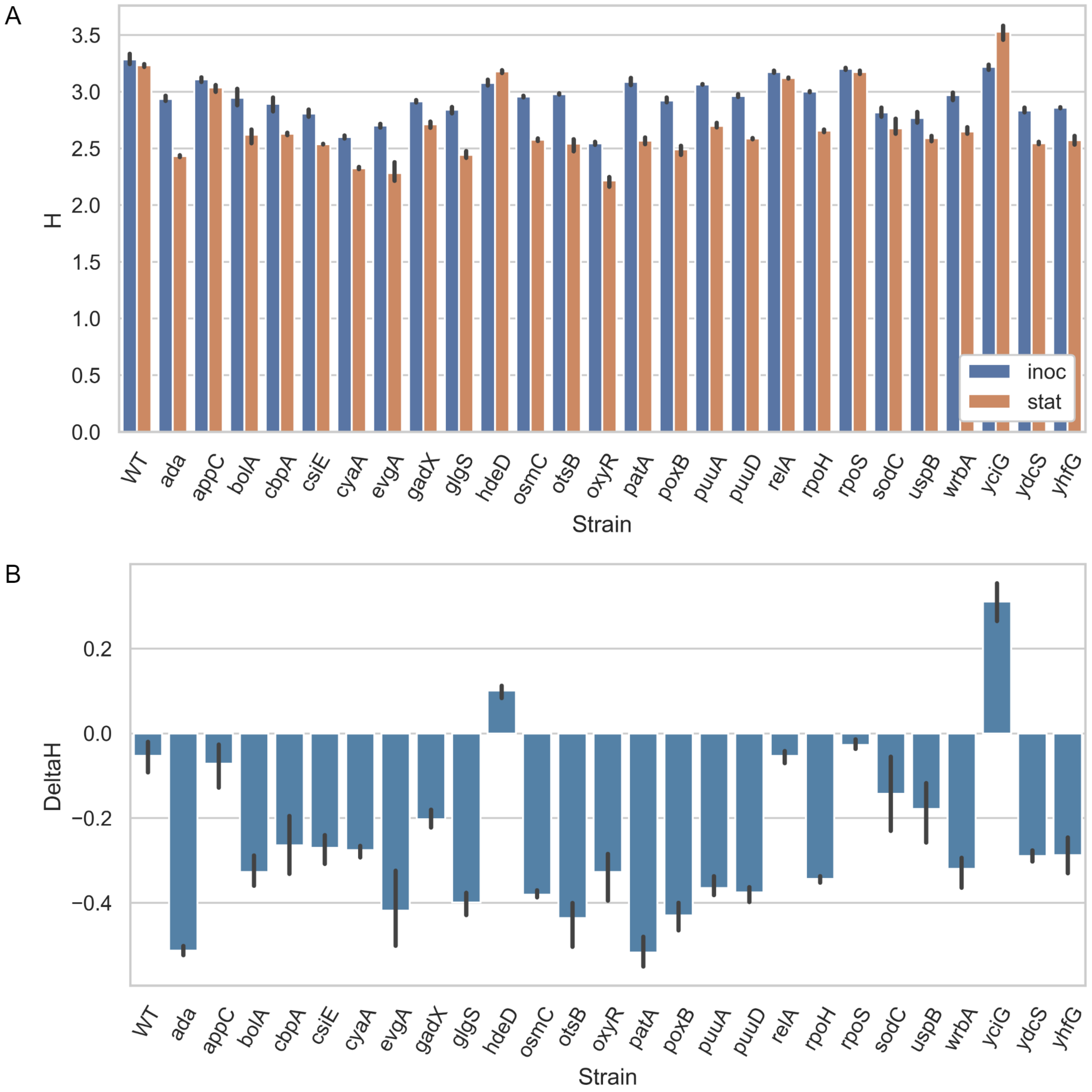

Figure S18: Shannon entropies of different transcriptional reporters sampled in the Biolector platform during exponential growth and stationary phase (**A**), and the differences of entropies between these phases (**B**).

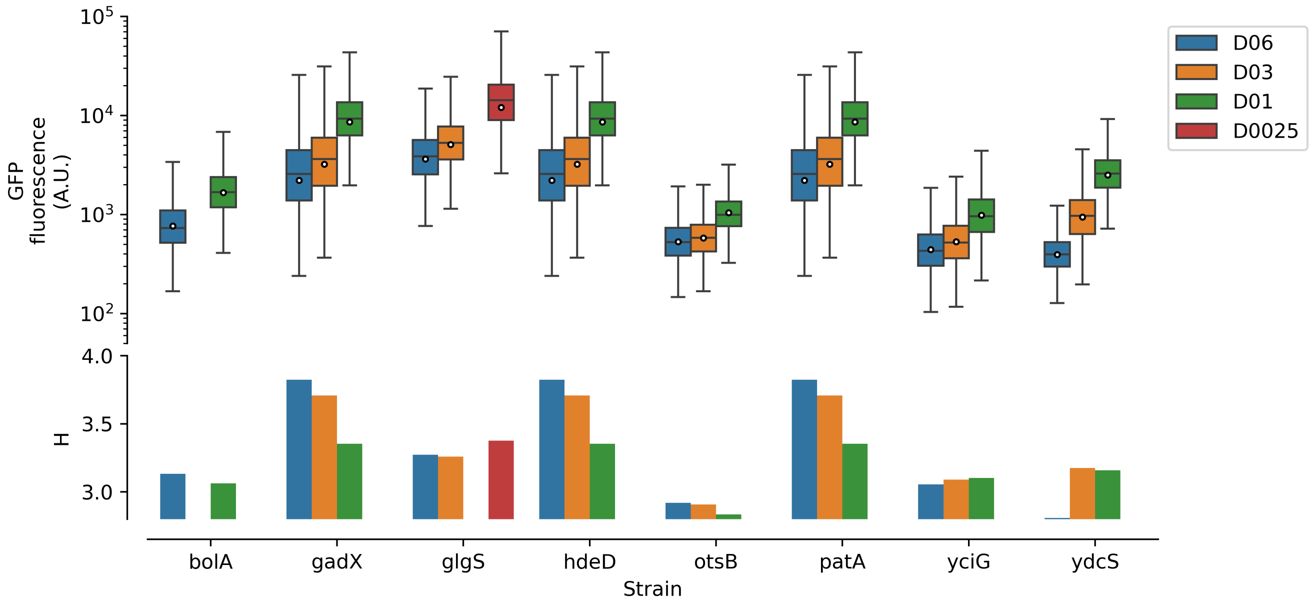

Figure S19: Summary plot of continuous cultures at different dilution rates performed with a subset of transcriptional reporters. The top panel represent the distribution of fluorescence as boxplots. The mean is indicated as a white dot and whiskers are drawn to the farthest datapoint within 1.5 * Interquartile range from the nearest hinge. The second panel represent the shannon entropies of corresponding samples.

#### 5.1. Extended materials and methods

##### 5.1.1. Strains and plasmids

*E. coli MG1655* strains from the Zaslaver collection [10] were used for the reporters screenings. Each strain carries a plasmid containing a transcriptional reporter (production of GFPmut2 controlled by the reported promoter) and confers kanamycin resistance. The promoters analyzed during the first screening in the Biolector platform are : $P_{ada}$, $P_{appC}$, $P_{bolA}$, $P_{cbpA}$, $P_{csiE}$, $P_{cyaA}$, $P_{evgA}$, $P_{gadX}$, $P_{glgS}$, $P_{hdeD}$, $P_{osmC}$, $P_{otsB}$, $P_{oxyR}$, $P_{patA}$, $P_{poxB}$, $P_{puuA}$, $P_{puuD}$, $P_{relA}$, $P_{rpoH}$, $P_{rpoS}$, $P_{sodC}$, $P_{uspB}$, $P_{wrbA}$, $P_{yciG}$, $P_{ydcS}$ and $P_{yhfG}$. In addition, a *E. coli MG1655* strain without plasmid has been used as control. The strains used for the second step of screening in continuous cultures were the ones with the following promoters: $P_{bolA}$, $P_{gadX}$, $P_{glgS}$, $P_{hdeD}$, $P_{otsB}$, $P_{patA}$, $P_{yciG}$ and $P_{ydcS}$. The dual reporter strain used in a Microfluidic Single Cell Cultivation device and the related experiments are described in the main document.

##### 5.1.2. Screening in Biolector platform

**Cultivation in the Biolector platform:** The first screening was performed in a BioLector (M2PLabs, Germany) with 48-deepwells microplates allowing measurement of Biomass (filter module 401: Exc. 620/10 nm | scattered light) and GFP (filter module 404: Exc 480/10 nm | Em. 520/25 nm). The mineral medium used was the same as described in the main document, with the addition of 10 g/L of MOPS as buffer (pH=7). Medium without kanamycin was used with the strain without plasmid. The strains were pre-cultivated in two steps to avoid initial induction of the reporters. The first precultures were inoculated in 50 ml flasks (with 5ml of the corresponding medium) with one single colony from a LB plate (supplemented with 50 mg/L kanamycin, for strains with plasmid). It was then incubated overnight at 37 °C at a shaking speed of 150 rpm. 8 µl of each of the first precultures were added to three different wells (triplicates) of black bottom biolector flower-plates (ref: MTP-48-OFF) containing 800 µl of fresh medium and were cultivated in Biolector (shaking frequency = 1400 rpm, Temperature = 37°C). After 4 hours, 8 µl of each of these second precultures were transfered in 800 µl of fresh medium in correspondings wells of readable biolector flower-plate (ref: MTP-48-B) and incubated in the same conditions. Three wells were used as blank and contained only medium. For each well, biomass and GFP were mesured every approximately 5.5 min. The remaining cell suspensions of the second precultures were analyzed in flow-cytometry as the sample for exponential growing phase (supposedly with the minimum fluorescence) and their $OD_{600}$ have been measured to approximate the initial $OD_{600}$ of the cultures. After more than 20 hours, when the culture reached stationary phase, 20 µl of each well was sampled for cytometry analysis. This volume was replaced by 20 µl of a concentrated glucose solution (200 g/L) and the plate was further incubated for about 20 hours. The data are available in the supplementary dataset and the code is available in the `Supp_Biolector_Process_Data.py` file.

**Off-line cytometry analysis:** Flow cytometry analyses were performed using a benchtop flow cytometer (BD Accuri C6) with the medium pre-set flow rate (35  L/min) and core sizes (16  m). The primary threshold was set at 20 000 units of FSC-A. The FL1-A channel (exc. 488 nm, em. 533/30 nm) was used for GFP quantification. Flow cytometry data was cleaned before being used by removing zeros and doublets. A second threshold was set at 10 000 units of FSC-A to correct an artefact in the measurements. To compute the entropy, 50 fixed bins in log scale between 0 and $10^6$ a.u. of GFP fluorescence were used. It is important to note that the values of entropies cannot be compared between different cytometers (e.g., BD Accuri C6 and BD Accuri C6+) and different fluidics parameters (i.e., flow-rates, core-size and thresholds). Since the other cultivations and the calibrations were not carried out with the exact same parameters, they cannot be compared to these measurements. The data are available in the supplementary dataset and the code is available in the `Supp_Biolector_FCA.py` file.

##### 5.1.3. Screening in chemostat experiments at three different dilution rates

**Cultivation in the Dasgip bioreactors:** Precultures were performed in two steps, as described in the main document. Chemostat experiments were performed in Dasgip bioreactors (Eppendorf, Germany) with 150 mL of working volume at controlled pH 7, 1000 rpm, and 37°C. Three dilution rates, i.e., 0.6, 0.1, and 0.3 h-1, were sequentially imposed on the system. Details about samplings and timings of experiments can be found in the supplementary document "`Supp_EcoliStressReporterSelection/Dasgip/Notes-MetaData.docx`".

**Online cytometry analysis:** For automated flow-cytometry analyses of 3 Dasgips reactors running in parallel, the “multi-stat” system was used. It is an adaptation of our in-house Segregostat [13] connected to a benchtop flow cytometer (BD Accuri C6). Every approximately 15 min, it takes a sample from one of the bioreactor and dilutes it with PBS before analysis. The following measure is performed in the next reactor, meaning that a sample is taken every approximately 45 min in each reactor. The fluidics flow rate was custom (24 µL/min, 8 µm core size) and a threshold was set either to 20 000 units of FSC-A. The FL1-A channel (exc. 488 nm, em. 533/30 nm) was used for GFP quantification. Flow cytometry data was cleaned before being used by removing zeros and doublets. To compute the entropy, 50 fixed bins in log scale between 0 and $10^6$ a.u. of GFP fluorescence were used. The cytometry files ('.fcs') used to make the Figure S19 correspond to the last measure in the last retention time for each dilution rate. They are listed in the supplementary document `"Supp_EcoliStressReporterSelection/Dasgip/Notes-MetaData.docx"`. The data are available in the supplementary dataset and the code is available in the `Supp_Dasgip_FCA.py` file.

### Supplementary Note 6. Time-lapse microscopy in a Microfluidic Single Cell Cultivation (MSCC) device

The dual-reporter strain *E. coli* MG1655 $P_{rpoS}$::CyOFP1 $P_{ydcS}$::GFPmut2 was characterized using time-lapse microscopy in a Microfluidic Single Cell Cultivation (MSCC) device (Figures S20-S22).

As described in the main text, the trade-off between growth and gene expression of *S.cerevisiae* $P_{glc3}$::GFP was also monitored in this device (Figure S23).

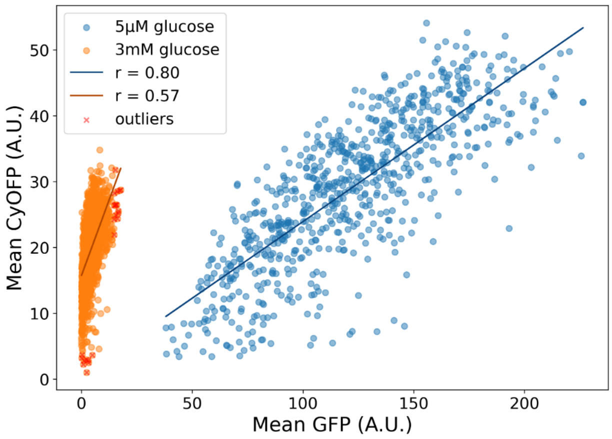

Figure S20: Correlations between $P_{rpoS}$::CyOFP1 and $P_{ydcS}$::GFPmut2 expression in limiting (5 µM glucose, blue) and non-limiting (4 mM glucose, orange) conditions in a MSCC device. Each dot represent a single-cell (from a total of 5 cultivation chambers per glucose concentration condition). Outliers are datapoint with a $|z\text{-score}| > 3$.

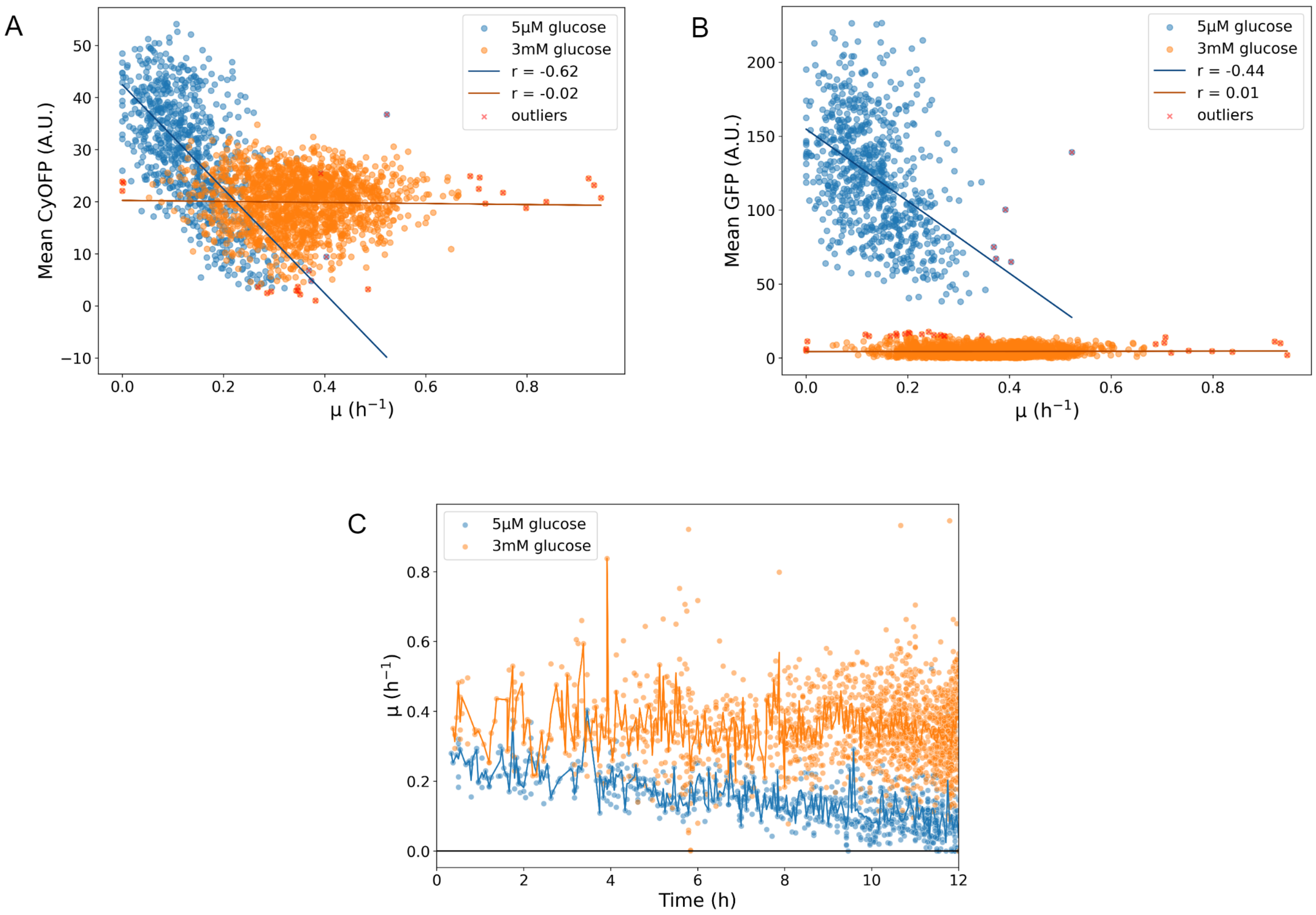

Figure S21: Single Cell growth in limiting (5 µM glucose, blue) and non-limiting (4 mM glucose, orange) conditions in a MSCC device. Each dot represent a single-cell (from a total of 5 cultivation chambers per glucose concentration condition). Outliers are datapoint with a $|z\text{-score}| > 3$. (A) Correlations between $P_{rpoS}$::CyOFP1 expression and growth rate, (B) Correlations between $P_{ydcS}$::GFPmut2 expression and growth rate, (C) Evolution of single-cell growth rate over time.

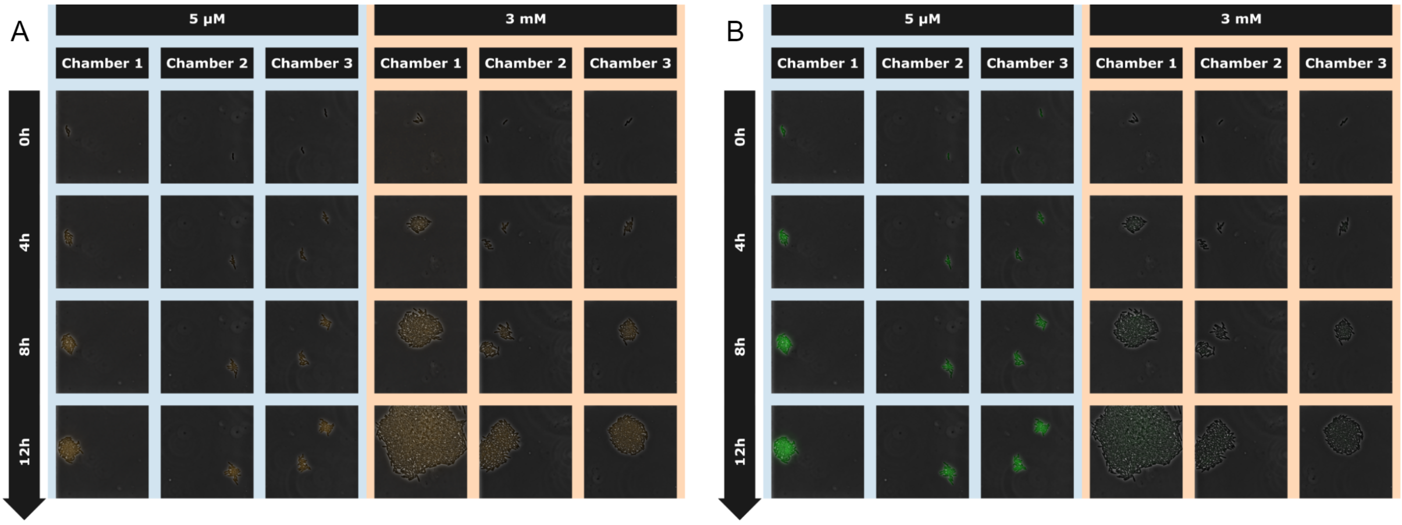

Figure S22: Time-lapse microscopy pictures of *E.coli* $P_{rpoS}$::CyOFP1 $P_{ydcS}$::GFPmut2 cultivated in MSCC,three microfluidic chambers per condition. Stressfull condition ([glu] = 5 µM) is indicated with blue background and non-stressful condition ([glu] = 3 mM) is indicated with orange background. (A) CyOFP fluorescence. (B) GFP fluorescence.

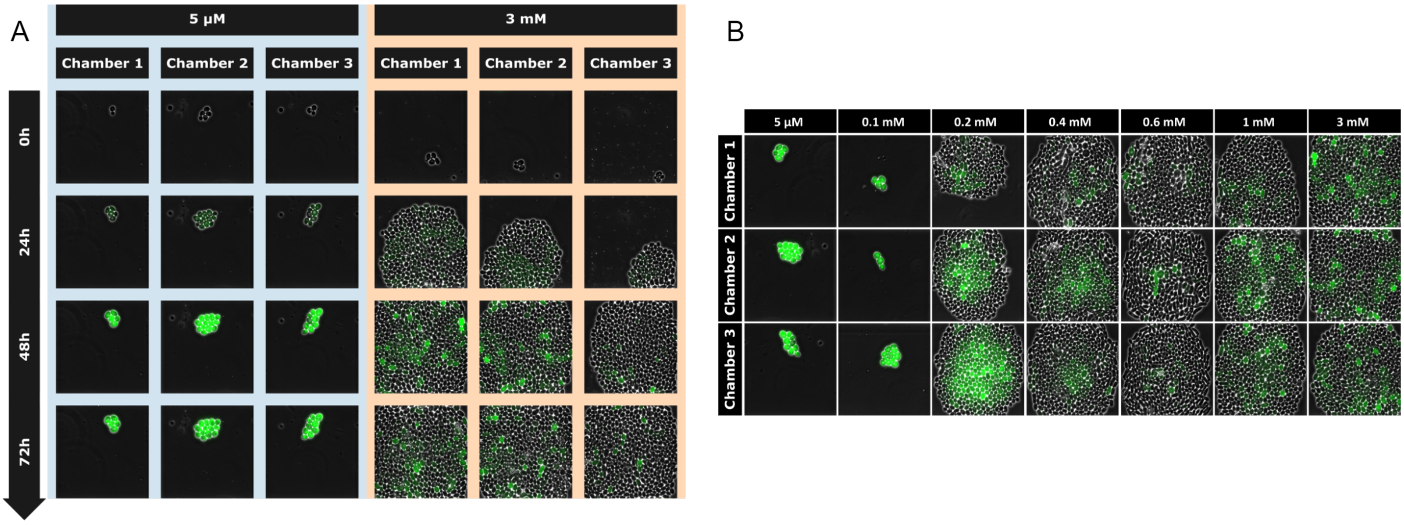

Figure S23: Microscopy pictures of *S.cerevisiae* $P_{glc3}$::GFP cultivated in MSCC, three microfluidic chambers per condition. (A) Time-lapse in stressfull condition ([glu] = 5 µM) indicated with blue background, and in non-stressful condition ([glu] = 3 mM), indicated with orange background. (B) Microscopy pictures at different GLU feed concentrations (0.005, 0.1, 0.2, 0.4, 0.6, 1 and 3 mM) after 48 h of cultivation (from Martinez et al. [14]).

### Supplementary Note 7. Cultivations of *S. cerevisiae* $P_{glc3}$::GFP at various dilution rates

Residual glucose concentrations at different dilution rates were obtained through changestat experiments [15]. In these continuous cultures, the feed pump speed was either increased (accelerostat,

A-stat) or decreased (decelerostat, D-stat) in steps of 1 rpm every two hours, resulting in a dilution rate change of 0.004 $^{-1}$ per step. This strategy was chosen to ensure a quasi-steady state at each step.

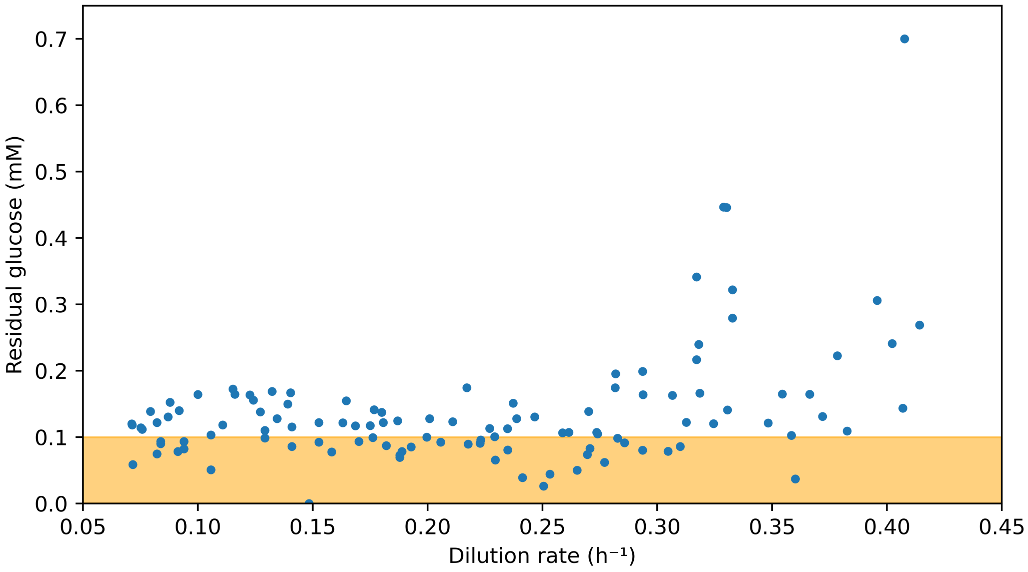

Figure S24: Residual glucose concentrations at different dilution rates (data from Maximilian Sehrt's work [15])

### 8. Tables

#### 8.1. Strains and plasmids

Note: the uppercase sequences in the pTarget primers are the selection markers for the CRISPR-cas9 system.

Table 1: The uppercase sequences in the pTarget primers are the selection markers for the CRISPR-cas9 system.

| Related Strain | Role | Primer name | Sequence (5'-3') |
|---|---|---|---|
| E.coli MG1655 *PrpoS*::CyOFP1 | Cassette construction | Fw Homo1 | cgcagagcaactggataagcc |
| | | Rv Homo1 + Tail-RBS | ctagtatttctcctctttctctagattactcgcggaacagcgc |
| | | Fw CyOFP1 + Tail-RBS | tctagagaaagaggagaaatactagatggtgagcaagggcgagg |
| | | Rv CyOFP1 | ttacttgtacagctcgtccatgcc |
| | | Fw Homo2 + tail-CyOFP1 | ggacgagctgtacaagtaagtaagcatctgtcagaaagg |
| | | Rv Homo2 | agccgcatttattatttcactgg |
| | Construction of pTarget | Fw pTarget-rpoS | agctagctcagtcctaggtataatactagt GACAGATGCTTACTTACTCG gttttagagctagaaatagcaagttaaaa |
| | | Rv pTarget | actagtattatacctaggactgagctagctg |
| | | Fw verif_pTarget-rpoS | gacagatgcttacttactcg |
| | | Rv verif_pTarget | ttgccgactaccttggtgat |
| | Verification/Sequencing | Fw verif_prpoS-CyOFP1 | tggttactggtgctggagg |
| | | Rv verif_prpoS-CyOFP1 | ccgtactattcgtttgccg |

Table 2: The uppercase sequences in the pTarget primers are the selection markers for the CRISPR-cas9 system. For the lacI transformation, selecting a lacI pTarget makes the pTarger and the pCas plasmid mutually exclusive. It is possible to select for the modification with the second antibiotic, but pCas is removed from the selected cells. Then, the pTarget can be cured by electroporating the pCas plasmid back in the strain.

| Related Strain | Role | Primer name | Sequence (5'-3') |
|---|---|---|---|
| E.coli MG1655 ΔlacAYZ ΔlacI | Cassette construction | Fw Homo1_lacAYZI | ccggtattgtgggcgatggt |
| | | Rv Homo1_lacAYZI | gagtcaattcagggtggtgaatattataaaaattgcctgatacgctgcg |
| | | Fw Homo2_lacAYZI | attcaccaccctgaattgactc |
| | | Rv Homo2_lacAYZI | ggtgatgactatcaactggcacgg |
| | Construction of pTarget | Fw pTarget-lacI | agctagctcagtcctaggtataatactagt AACGGTCTGATAAGAGACAC gttttagagctagaaatagcaagttaaaa |
| | | Rv pTarget | actagtattatacctaggactgagctagctg |
| | | Fw verif_pTarget-lacI | AACGGTCTGATAAGAGACAC |
| | | Rv verif_pTarget | ttgccgactaccttggtgat |
| | Verification/Sequencing | Fw verif_lacAYZI | tatggctcgccatcaggatcgg |
| | | Rv verif_lacAYZI | ttatcctttcaccgggcaatggt |
| E.coli MG1655 ΔlacAYZ ΔlacI ΔserA | Cassette construction | Fw Homo1_serA | gtggtgaccgataatggcaacgtgat |
| | | Rv Homo1_serA | ttacccaatcctgtcttttgaaatgttgtgtgc |
| | | Fw Homo2_serA | gacaggattgggtaattccccttctctgaaaatcaacgggc |
| | | Rv Homo2_serA | ctgcggtggttacaccgtcgaa |
| | Construction of pTarget | Fw pTarget-serA | agctagctcagtcctaggtataatactagt AGACAAGATTAAGTTTCTGC gttttagagctagaaatagcaagttaaaa |
| | | Rv pTarget | actagtattatacctaggactgagctagctg |
| | | Fw verif_pTarget-serA | agacaagattaagtttctgc |
| | | Rv verif_pTarget | ttgccgactaccttggtgat |
| | Verification/Sequencing | Fw verif_serA | gtattgcagacgcttccaagcagg |
| | | Rv verif_serA | ccgattagacgtgctgatcacacc |

#### 8.2. Lab-scale bioreactors cultivations

Table 3: Summary of specific operating conditions of the continuous cultures performed in lab-scale bioreactors.

| ***Culture type and strain*** | ***Operating conditions*** | ***Replicates*** | ***Publication*** |
|---|---|---|---|
| Segregostat<br>*E. coli* W3110 $P_{araB}$::GFPmut2 | Bioreactor : Biostat B-Twin (Sartorius)<br>Cytometer : BD Accuri C6 (BD Biosciences)<br>D = 0.45 $h^{-1}$ pH = 7 T = 37 °C<br>Aeration = 1 L/min Agitation = 1000 rpm<br>Regulation threshold = 1000 A.F.U.<br>$[glucose]_{FEED}$ = 5 g/L<br>Regulation pulse = 0.15 g arabinose | 2021-09-29_LHE_Seg_Ecoli_Ara<br>2022-01-05_LHE_Seg_Ecoli_Ara | Henrion et al. (2023) |
| Segregostat<br>*E. coli* BL21(DE3) pET28::GFP | Bioreactor : Biostat B-Twin (Sartorius)<br>Cytometer : BD Accuri C6 (BD Biosciences)<br>D = 0.45 $h^{-1}$ pH = 7 T = 37 °C<br>Aeration = 1 L/min Agitation = 1000 rpm<br>Regulation threshold = 1000 A.F.U.<br>$[glucose]_{FEED}$ = 5 g/L<br>Regulation pulse = 0.15 g lactose | 2022-09-20_LHE_Seg_Ecoli_BL21<br>2022-10-11_LHE_Seg_Ecoli_BL21 | Henrion et al. (2023) |
| Chemostat<br>*E. coli* BL21(DE3) pET28::GFP | Bioreactor : Biostat B-Twin (Sartorius)<br>Cytometer : BD Accuri C6 (BD Biosciences)<br>D = 0.45 $h^{-1}$ pH = 7 T = 37 °C<br>Aeration = 1 L/min Agitation = 1000 rpm<br>$[glucose]_{FEED}$ = 5 g/L<br>$[lactose]_{FEED}$ = 1 g/L | 2022-08-10_LHE_Chem_Ecoli_BL21 | Henrion et al. (2023) |
| | Bioreactor : Bionet F1 bioreactor (Bionet)<br>Cytometer : BD Accuri C6+ (BD Biosciences)<br>D = 0.45 $h^{-1}$ pH = 7 T = 37 °C<br>Aeration = 1 L/min Agitation = 1000 rpm<br>$[glucose]_{FEED}$ = 5 g/L<br>$[lactose]_{FEED}$ = 1 g/L | 2023-02-15_LHE_Chem_Ecoli_BL21 | Henrion et al. (2023) |
| | | 2025-03-11_ALS_Chem_Ecoli_BL21<br>2025-04-01_ALS_Chem_Ecoli_BL21 | This paper |
| Chemostat xylose<br>*E. coli* BL21(DE3) pET28::GFP | Bioreactor : Bionet F1 bioreactor (Bionet)<br>Cytometer : BD Accuri C6+ (BD Biosciences)<br>D = 0.45 $h^{-1}$ pH = 7 T = 37 °C<br>Aeration = 1 L/min Agitation = 1000 rpm<br>$[xylose]_{FEED}$ = 5 g/L<br>$[lactose]_{FEED}$ = 1 g/L | 2025-05-05_ALS_Chem_Ecoli_BL21_xylose<br>2025-05-28_FDE_Chem_Ecoli_BL21_xylose | This paper |
| Chemostat<br>*E. coli* MG1655 ΔlacAYZ ΔlacI pECJ3 | Bioreactor : Bionet F1 bioreactor (Bionet)<br>Cytometer : BD Accuri C6+ (BD Biosciences)<br>D = 0.5 $h^{-1}$ pH = 7 T = 37 °C<br>Aeration = 1 L/min Agitation = 1000 rpm<br>$[glucose]_{FEED}$ = 5 g/L<br>Induction : after 12h of continuous culture<br>Inducer : medium with 0.1 mM IPTG | 2024-02-14_VIV_ToggleControl1<br>2024-05-09_VIV_ToggleControl2 | This paper |
| Chemostat<br>*E. coli* MG1655 ΔlacAYZ ΔlacI ΔserA pECJ3 (PLtetO-1::serA::GFP) | | 2024-03-13_VIV_ToggleTSG1<br>2024-03-27_VIV_ToggleTSG2 | This paper |
| Segregostat<br>*E. coli* MG1655 $P_{ydcS}$::GFPmut2 | Bioreactor : Bionet F1 bioreactor (Bionet)<br>Cytometer : BD Accuri C6+ (BD Biosciences)<br>D = 0.3 $h^{-1}$ pH = 7 T = 37 °C<br>Aeration = 1 L/min<br>Agitation : 1000 & 1200 rpm<br>$[glucose]_{FEED}$ = 5 g/L<br>Regulation threshold = 1000 & 1150 A.F.U.<br>Regulation pulse = 1 g glucose (5.7 ml) | 2024-09-11_FDE_Seg_Ecoli_ydcS<br>2025-01-29_MaD_Seg_Ecoli_ydcS | This paper |
| *Segregostat*<br>*S. cerevisiae* CEN.PK 113-7D $P_{glc3}$::eGFP | Bioreactor : Biostat B-Twin (Sartorius)<br>Cytometer : BD Accuri C6 (BD Biosciences)<br>D = 0.1 $h^{-1}$ pH = 5 T = 30°C<br>Aeration = 1 L/min Agitation : 1000 rpm<br>$[glucose]_{FEED}$ = 5 g/L<br>Regulation threshold = 5000 A.F.U.<br>Regulation pulse = 0.4 g glucose | 2020-11-09_FDE_Seg_Sacch<br>2021-04-15_FDE_Seg_Sacch | Henrion et al. (2023) |
| | Bioreactor : Bionet F1 bioreactor (Bionet)<br>Cytometer : BD Accuri C6+ (BD Biosciences)<br>D = 0.1 $h^{-1}$ pH = 5 T = 30°C<br>Aeration = 1 L/min Agitation : 1000 rpm<br>$[glucose]_{FEED}$ = 5 g/L<br>Regulation threshold = 3000 A.F.U.<br>Regulation pulse = 0.4 g glucose (5.7 ml) | 2025-02-10_MaD_Seg_Sacch | This paper |
| *Chemostat*<br>*E. coli MG1655 PydcS::GFPmut2* | Bioreactor : Bionet F1 bioreactor (Bionet)<br>Cytometer : BD Accuri C6+ (BD Biosciences)<br>D = 0.6 & 0.1 $h^{-1}$ pH = 7 T = 37 °C<br>Aeration = 1 L/min Agitation = 1000 rpm<br>$[glucose]_{FEED}$ = 5 g/L | 2024-06-12_MaD-FDE_Chem_Ecoli_ydcS<br>2024-06-26_MaD_Chem_Ecoli_ydcS | This paper |
| | Bioreactor : Bionet F1 bioreactor (Bionet)<br>Cytometer : BD Accuri C6+ (BD Biosciences)<br>D = 0.3 $h^{-1}$ pH = 7 T = 37 °C<br>Aeration = 1 L/min Agitation : 1000<br>$[glucose]_{FEED}$ = 5 g/L | 2025-01-29_MaD_Chem_Ecoli_ydcS | This paper |
| | Bioreactor : Dasgip (Eppendorf, Germany)<br>Cytometer : BD Accuri C6 (BD Biosciences)<br>D = 0.6, 0.3 & 0.1 $h^{-1}$ pH = 7 T = 37 °C<br>Aeration = 1 L/min Agitation : 1000<br>$[glucose]_{FEED}$ = 5 g/L | See the "Supp_EcoliStressReporterSelection" folder in provided data | This paper |
| *Chemostat*<br>*S. cerevisiae* CEN.PK 113-7D $P_{glc3}$::eGFP | Bioreactor : Bionet F1 bioreactor (Bionet)<br>Cytometer : BD Accuri C6+ (BD Biosciences)<br>D = 0.3 & 0.1 $h^{-1}$ pH = 5 T = 30°C<br>Aeration = 1 L/min Agitation : 1000 rpm<br>$[glucose]_{FEED}$ = 7.5 g/L | 2024-07-04_MaD_Chem_Sacch<br>2024-08-19_MaD_Chem_Sacch | This paper |

[3] Georges Vicqueray. "Characterisation and Modelisation of Phenotypic Variability in Saccharomyces Cerevisiae: The Interaction of Nutrient Stress, Cell Size, and Morphology." MA thesis. University of Liège, Aug. 2023.

[15] Maximilian Sehrt (Tieke). “Using Substrate Fluctuations For Controlling Bet-Hedging In Yeast Based On Automated Single-Cell Technologies”. MA thesis. Bielefeld, June 2022.
